# The Role of Glycosphingolipids in Autoimmune Manifestations and Myeloma in Gaucher Disease

**DOI:** 10.1101/2025.07.28.666118

**Authors:** E.V. Pavlova, J. Archer, M. Guimaraes Da Lomba Ferraz, J.M.F.G Aerts, T.M Cox, the MRC Gaucherite consortium

**Author notes:** Address correspondence to: Elena V. Pavlova, Box 157 Department of Medicine, University of Cambridge, Addenbrooke’s Hospital, Cambridge, CB2 0QQ, UK.

## Abstract

Gaucher disease, an inborn error of glucosylceramide recycling, predisposes to haematological malignancies, liver cirrhosis and cancer. Inducible Gaucher disease modelled in susceptible mouse strains showed striking autoimmune hepatitis and cancer development; as reported previously, they also develop monoclonal immunoglobulins and B-cell tumours. Exposure to eliglustat, a therapeutic inhibitor of glucosylceramide biosynthesis, suppressed hepatic disease and occurrence of monoclonal immunoglobulins. To interrogate activation of CD1-restricted T cells by glucosylceramides, we deleted CD1d1/Cd1d2 function in the conditional inducible murine model of Gaucher disease with GBA1 deficiency in haematopoietic cells. Systemic inflammation and autoantibodies in these CD1^-/-^ GCflox/flox Cre+ mice were markedly reduced, and severe liver injury and tumour formation was suppressed; however, occurrence of monoclonal immunoglobulins was unaffected. We suggest that CD1 molecules have distinct actions on tissue homeostasis in this system related to cell-intrinsic activation as well as glycolipid loading and presentation. Our findings implicate glycosphingolipids in the development of autoimmunity and cancer in human Gaucher disease and have broader significance in acquired and genetic disorders affecting sphingolipid expression.

## Introduction

Gaucher disease (GD), an inborn error of sphingolipid metabolism, is caused by biallelic mutations in *GBA1* that encodes lysosomal acid β-D-glucosylceramidase (Rosenbloom and Weinreb, 2013; Stirnemann et al., 2017). The condition is highly pleiomorphic but principally affects the bone marrow, liver and spleen which are infiltrated by alternatively activated macrophages (Boven et al., 2004); these are engorged with glucosylceramides and their deacylated derivative, glucosylsphingosine.

Impaired glycosphingolipid recycling in Gaucher disease provides a framework for exploring the role of sphingolipids in chronic inflammation associated with self-reactive B and T cell stimulation, autoantibody production and cancer immunology. The disease is notable for a greatly increased frequency of B cell lymphoma, myeloma and occurrence of hepatocellular carcinoma (Ayto and Hughes, 2013; de Fost et al., 2008; Mistry et al., 2013; Regenboog et al., 2018; Rosenbloom et al., 2022; Weinreb et al., 2018; Weinreb and Lee, 2013). Polyclonal and monoclonal gammopathy are frequent in patients and occur also in experimental Gaucher disease modelled in mice (de Fost et al., 2008; Arends et al., 2013; Pavlova et al., 2013).

The clonal plasma-cell condition Monoclonal Gammopathy of Undetermined Significance (MGUS) occurs in 3% of the general population over 50 and in 5% of those over 70 years. MGUS is characterized by circulating monoclonal (M)-proteins and leads to the second most common hematologic malignancy, multiple myeloma with ≈1% annualised risk (Kyle et al., 2018).

We reported clonal expansion of B cells harbouring immunoglobulin gene rearrangements which progressed to B cell lymphoma or myeloma in an inducible model of murine Gaucher disease (Pavlova et al., 2013). We later demonstrated that eliglustat, a potent inhibitor of glucosylceramide biosynthesis (McEachern et al., 2007) and a therapy approved worldwide for Gaucher disease (Mistry et al 2024), suppressed lymphocyte proliferation, decreased polyclonal IgG, and prevented the appearance of paraproteins (monoclonal gammopathy) as well as B cell lymphoma and myeloma (Pavlova et al., 2015). The strong preventative effect, linked to reduced pathologic glycosphingolipids, immediately implicated endogenous glucosylceramides and glucosylsphingosine as driving factors in the development of cancer, B cell proliferation and systemic inflammation.

The clonal immunoglobulins occurring in patients with Gaucher disease have been reported to bind to glucosylsphingosine, suggesting antigen-driven selection in the expansion of B cells after presentation by CD1d-restricted human and murine type II glucosylsphingosine -specific natural killer T (NKT) cells constitutively expressing the T-follicular helper 2 phenotypes (Nair et al., 2015). Moreover, expansion of B-cells that secrete monoclonal antibodies which specifically bind glucosylsphingosine was postulated by these authors to trigger development of myeloma in patients with Gaucher disease; the antibodies were also found in about one third of persons with sporadic monoclonal gammopathies (Nair et al 2016).

Here we report studies in chimeric F1 mixed background Gaucher mice in which glycosphingolipid accumulation led to sustained immune activation with autoimmunity, as well as B cell lymphoma. Pathognomonic histological and immunological features of severe chronic active hepatitis were present and with ageing, several animals developed frank cirrhosis and liver tumours (hepatocellular carcinoma and cholangiocarcinoma). These findings mirror hitherto unexplained aspects of liver disease and cancer in Gaucher patients (Lachmann et al 2000; Regenboog et al, 2018; Rosenbloom et al., 2022).

Since the CD1d family of transmembrane glycoproteins related to MHC Class I molecules present autochthonous lipid antigens at the surface of antigen-presenting cells, to explore the putative role of murine CD1d and cognate Gaucher lipids glucosylceramide and glucosylsphingosine in the pathogenesis of B cell proliferation and myeloma and novel observation of the autoimmune liver disease, we generated Gaucher mice lacking CD1d and administered eliglustat specifically to inhibit glycosphingolipid biosynthesis. Long-term exposure to eliglustat significantly reduced hepatic inflammation and prevented progression to B cell lymphoma/myeloma or other tumours in Gaucher mice, thereby confirming the pathological role of the excess glycosphingolipids. While systemic inflammation and autoantibodies were markedly reduced and severe liver injury and tumour formation was suppressed in genetically modified Gaucher mice lacking CD1d, irrespective of eliglustat treatment, monoclonal gammopathy occurred.

Accordingly, we examined sera obtained from patients with Gaucher disease in the pre- or early treatment period and in those with evidence of hepatic cirrhosis identified an unexpected frequency of tissue-specific autoantibodies. In patients with severe hepatic involvement and Gaucher disease, including cirrhosis, the pathologic glycosphingolipid abnormalities due to GBA deficiency were associated with tissue-specific autoimmunity.

## Results

### Gaucher mice develop autoimmune liver disease and systemic inflammation

We monitored 50 Gaucher and 28 strain control mice. Breeding 129/sv.B6 GD mice with the congenic B6 strain produced offspring that developed an exacerbated autoimmune phenotype, which was triggered after induction of Gaucher disease and the accumulation of glycosphingolipids. As previously observed (Pavlova et al., 2013a, 2015a) 13 out of 50 Gaucher mice (26%) had diffuse large B cell lymphomas. Several GD mice after 11 months rapidly developed signs of illness, distended abdomen, weight reduction and were killed (Fig. 1). Histological examination revealed systemic pathology affecting several organs including liver, lungs, intestine, mesentery and kidneys. Infiltration by lymphocytes and plasma cells accompanied by raised serum immunoglobulins was noted in liver, kidneys, lungs and intestine as previously observed in the original colony of Gaucher mice. Splenomegaly and disruption of splenic architecture, expansion of the red pulp and significant extramedullary haematopoiesis were present. Notably, infiltration with large multinucleated macrophages (Gaucher cells) occurred in spleen, liver, bone marrow and thymus.

**Figure 1.**
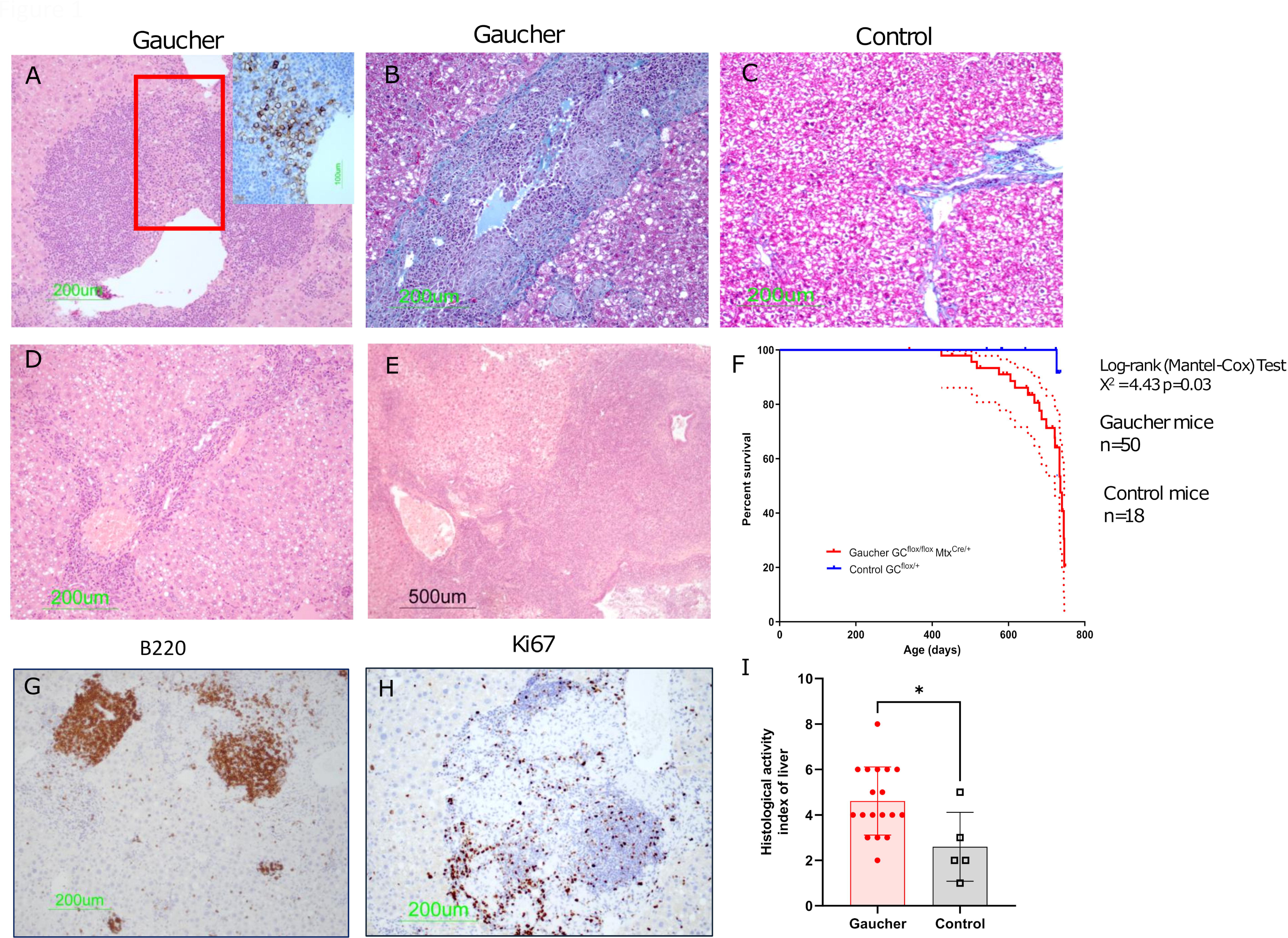
Histological features of autoimmune liver disease in. chimeric F1 mixed 129sv/B6 background Gaucher mice. A) Liver sections from Gaucher mouse with infiltrates of 138^+^ plasma cells. B) Liver fibrosis and collagen deposition stained with Masson Trichrome staining and (C) Control liver section of GC^flox/+^ mouse showing normal staining; D) haematoxylin & eosin stained liver section of Gaucher mouse showing increased ductal structures in the hepatic parenchyma; E) section with cholangiocarcinoma contained a large round structure rimmed by a single rim epithelium. In the dilated lumen necrotic material; F) Cancer incidence in 129/sv. B6 Gaucher strain (n=50) and control (n=18). Kaplan-Meier analysis censored at 24 months; G) B220^+^ B cells in liver section of Gaucher mouse showing infiltrates of B cells in the hepatic parenchyma; H) Ki-67^+^ proliferative cells in the hepatic infiltrate co-localised with enlarged Kupfer cells surrounded by mixed population of lymphocytes; I) Histological activity index of liver in Gaucher and control mice, bars and whiskers represent Mean and SD, *p<0.05.

Induced Gaucher mice developed autoimmune manifestations accompanied by lymphocytic infiltration of several organs but predominantly in the liver; in a proportion of these mice, florid lymphocytic infiltrates were present in the lungs, kidneys and intestine. Fifteen of 50 Gaucher mice (30%) had histological signs of hepatitis accompanied by cirrhosis, two animals had hepatocellular carcinoma, and one had cholangiocarcinoma; in one GD mouse, adenocarcinoma of the gallbladder was detected. There were multifocal areas of neutrophils, lymphocytes and macrophages present mainly in sinusoids, pericentral veins and portal areas. Large areas of necrosis with disrupted architecture, formation of irregular hepatic cords and nodules, and hepatocytes with significant vacuolation of cytoplasm were seen. Increased plasma cell infiltration characteristic of autoimmune hepatitis (Fig. 1A) and cholangitis were found in Gaucher mice (Fig. 1D, E). Elevation of alanine transaminase and glutamate dehydrogenase in GD mice with histological signs of hepatitis as well as modest increase of total bilirubin (Fig. 2C). Histological examination of several organs identified systemic inflammation in Gaucher mice resembling autoimmune disease.

**Figure 2.**
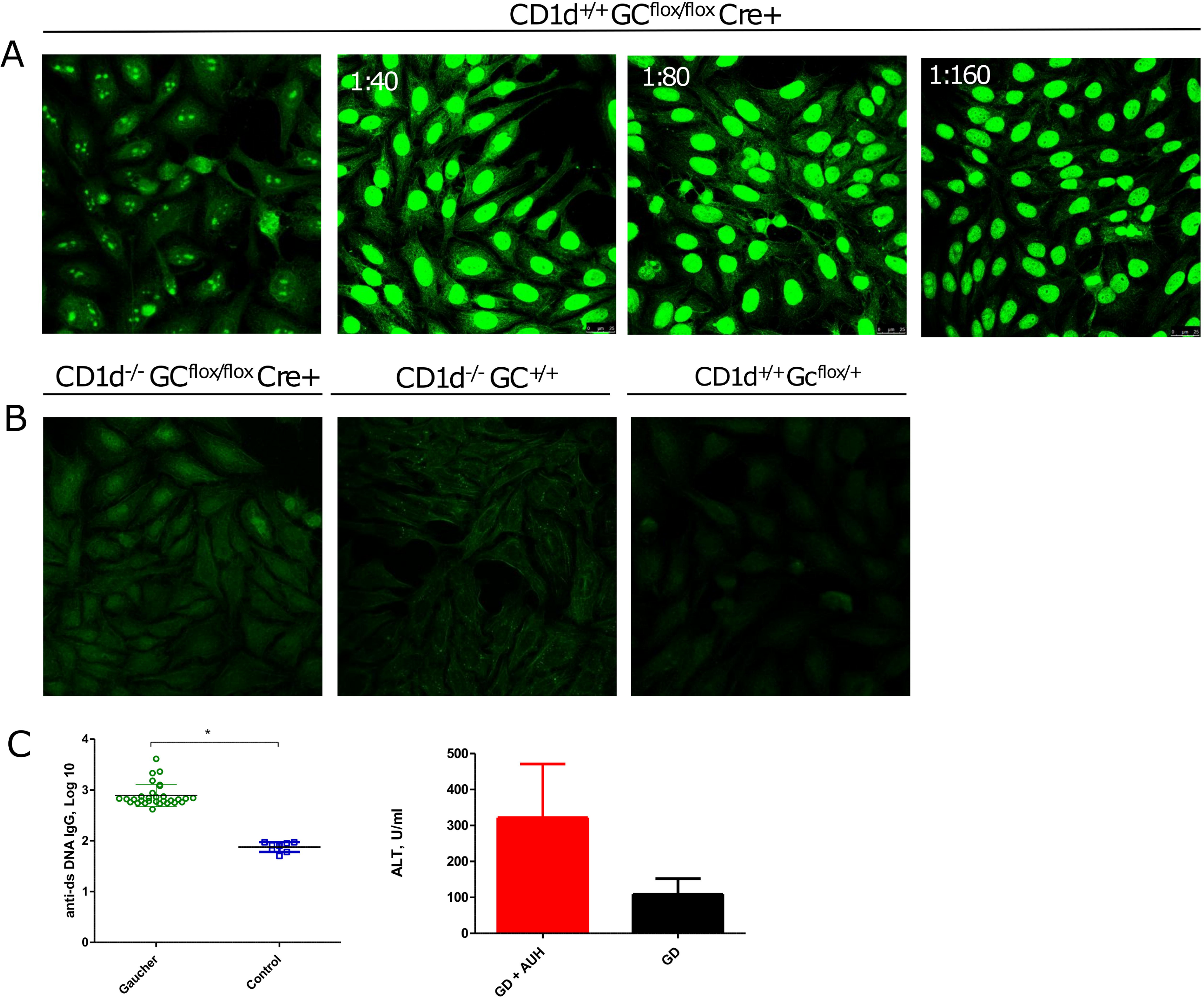
Morphologic characteristic of patterns of anti-nuclear autoantibodies. identified by indirect immunofluorescence on Hep-2 cells stained with serum from Gaucher and CD1d^-/-^ GC^flox.flox^ Cre+ mice: A) antinuclear serial dilution in Gaucher mice, B) Negative staining with serum from CD1d^-/-^ GC^flox.flox^ Cre+ mice; C) raised concentration of anti-dsDNA IgG and ALT in mice with autoimmune hepatitis. * p<0.05. Bars and whiskers represent Mean and SD.

Autoimmune hepatitis with systemic inflammation was triggered by the interferon stimulus and glycosphingolipid accumulation in Gaucher mice on a chimeric mixed background. Breakdown of peripheral tolerance may be attributed to abnormalities in B cell differentiation in chronic antigenic stimulation by endogenous glycosphingolipids combined with interferon-induced activation of antigen-presenting cells. The liver changes had the pathognomonic histological and immunological features of chronic active hepatitis, a condition due to impaired tissue self-tolerance. Hepatitis was moreover absent in animals with genotypes that do not allow generation of the Gaucher-related sphingolipid disease (eg. Gba flx +/-, Cre -ve). As required for a high-level animal facility, all mouse colonies were free of murine hepatitis virus.

### Autoantibodies ANA and anti-ds DNA in Gaucher mice

To test if systemic inflammation observed in experimental Gaucher mice was characteristic of autoimmune disease, we sought the presence of autoantibodies in GD mouse sera. Human epithelial type 2 (Hep-2) cells were stained with sera of GD and control mice. To search for autoantibodies in human autoimmune diseases that react characteristically with specific antigens, we tested for anti-mitochondrial, anti-ds-DNA, anti-ss DNA IgG in sera obtained from mice with Gaucher disease. Anti-nuclear antibody with speckled, nucleolar and anti-mitochondrial patterns was found in 9 out of 20 (45%) of Gaucher mice with elevated titres of anti-dsDNA IgG. (Fig. 2).

### Generation of CD1d deficiency in Gaucher mice

To investigate if CD1-dependent presentation of lipid self-antigens such as β-D-glucosylceramide and related glucosylsphingosines specific to Gaucher disease, drive antigen-mediated immune response and pre-malignant and malignant lymphocyte proliferation in the glycosphingolipid environment caused by glucosylceramidase deficiency, we crossed B6.129S2.BALB/c-Cd1^tm1Gru^ mice (Smiley et al., 1997) with Gba1^tm1Karl^/Gba1^tm1.1Karl^ <u>Tg(Mx1-cre)1Cgn</u>/0 (Gaucher) mice (Enquist et al., 2006). Murine CD1 antigen complex is encoded by two homologous *Cd1d1* and *Cd1d2* genes, that are located on mouse chromosome 3 in proximity of ∼2.8Mb to the *Gba1* gene (Fig. 3 A).

**Figure 3.**
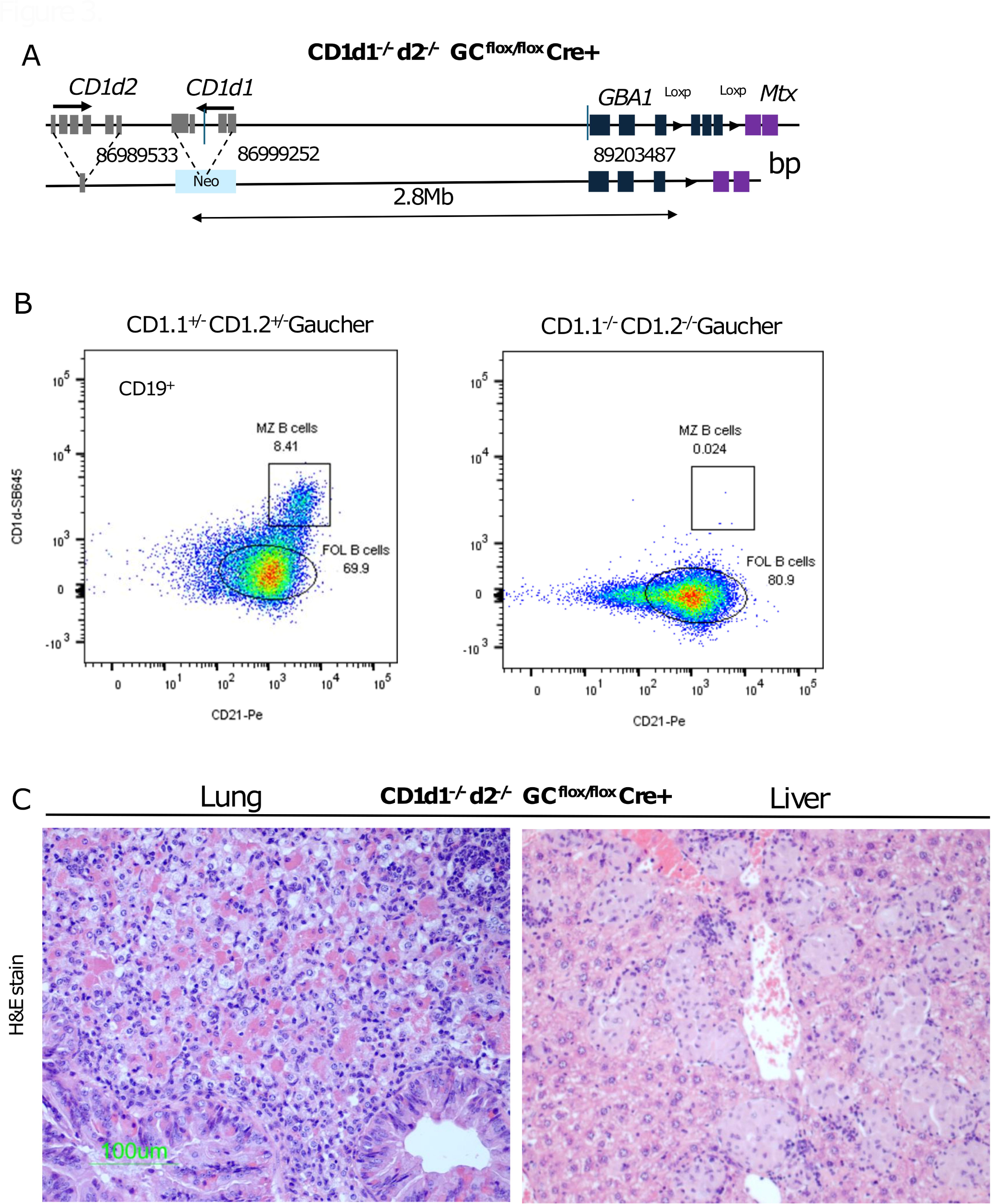
Generation of CD1d1-/-d2-/-GCflox/flox Cre+ mice. A) The schematic diagram of the genetic recombination involving *CD1* and *GBA1* targeting constructs on mouse chromosome 3qF1. The exons of the *CDd1.1, CDd1.2* and *GBA1* genes are shown. The resulting recombinant with neomycin insertion in the CD1d1, and LoxP insertions in the GBA1 loci. The physical distance between the CD1 and GBA1 loci is indicated. The locations based on NCBI mouse genome build GRCm38/mm10. B) Expression of CD1d in the spleen marginal zone B cells CD19^+^ CD21^+^ of a CD1d^+^Gaucher mouse and lack of CD1d expression in Cd1d1-/-d2-/-GCflox/flox Cre+, proportions of follicular B cells are also indicated; C) H&E stained section of the lung of a CD1d1-/-d2-/-GCflox/flox Cre+ mouse containing clusters of large macrophages with fibrillary eosinophilic material in the cytoplasm and infiltration of neutrophils and eosinophils. These cells surrounded bronchi and blood vessels. C) Multifocal collections of Gaucher’s cells and small lymphocytes in the liver parenchyma.

The first seven recombinant progeny were born among 66 pups (supplementary table TS1) due to an increased frequency of gametic crossovers between *Cd1* and *Gba1* genes which suggest a potential meiotic recombination hotspot site between these two loci. For the physical distance between the *Cd1* and *GBA1* loci 2.8Mb, estimated genetic interval would be ∼1.5cM in the mouse genome, where one centimorgan is approximately equivalent to 2,000 kilobases (kb). The recombinant male mouse harbouring Cd1 ^tm1Gru^ and Gba1^tm1Karl^ alleles did not produce offspring despite mating with multiple females. Examination revealed an absence of sperm in the seminal ducts, confirming sterility.

Subsequent sib-mating resulted in inbreeding depression in the Gaucher strain which was evidenced by lower fertility, reduced litter sizes and growth rates. When we later interbred the offspring of the sibling-mated GC^flox/+^ parents with CD1d+/-GCflox/+ mice, the second recombination event between the *CD1d* and *GBA1* loci was found to have occurred in the first filial generation (F1) hybrids CD1^+/-^ GC^flox/flox^. This phenomenon can be attributed to heterosis, resulting from the evolutionary advantage of the hybrids.

The F2 generations of the CD1d1-/-d2-/-GC^flox/flox^ mice appeared healthy and produced offspring at the expected Mendelian ratios. Mouse Universal Genotyping Array (Transnetyx) based on over 11,000 single nucleotide polymorphism (SNP) probes, identified informative SNPs indicating the presence of the background strain groups BALB/c, C57BL/6 and the substrains: BALB/cJ, C57BL/6J, C57BL/6JBomTac in our colony (Sigmon et al., 2020).

### Histological features in CD1d1-/-d2-/-GCflox/flox mice

Histological examination showed abundant storage macrophages in spleen, liver, lungs and thymus of CD1d^-/-^GC^flox/flox^ Cre+ mice (Fig. 3C). Parenchymal infiltration by mixed clusters of lymphocytes and plasma cells was noticeable in the lungs, liver, spleen and kidneys. Clusters of large macrophages containing fibrillary eosinophilic material in the cytoplasm were also scattered throughout, while a few alveolar macrophages were present. Mixed with these there were clusters of neutrophils and eosinophils surrounding bronchi and blood vessels. Multifocal collections of Gaucher cells along with variable numbers of small lymphocytes were found throughout the liver parenchyma. The lobes were formed from hepatocytes with granular mildly vacuolated cytoplasm. Perivascular and periportal small collections of small lymphocytes were widely distributed; multifocal mostly intra-sinusoidal round collections of Gaucher cells were also found.

In the kidneys, small lymphocytes and plasma cells were mostly in the pelvic area. Glomerular tufts were replaced by plasma cell and one was completely effaced, indicating glomerulonephritis. A few dilated tubules containing eosinophilic material were present. Scattered perivascular collections of small lymphocytes were found and large collections in occurred in the pelvis.

In the lungs the alveolar walls were mildly thickened, and the alveoli were irregular in size and shape; most had a clear lumen. Bronchi were intact with clear, dilated lumina. Blood vessels were intact and dilated with blood. Many perivascular and peribronchial collections, principally of small lymphocytes with scattered reactive plasma cells, were present.

Overall, the lupus-like histological features in CD1^-/-^ GC^flox/flox^ Cre+ mice involved the liver, kidneys and lungs with infiltration of plasma cells and increased histological activity resembling chronic inflammation but neither progressed to severe hepatic cirrhosis nor led to formation of any tumours.

### Monoclonal gammopathy and immunoglobulin concentrations in CD1^-/-^ GC^flox/flox^ Cre+ mice

High concentrations of glucosylceramide and glucosylsphingosine in thymus and spleen were associated with increased frequency of monoclonal immunoglobulins in Cd1d1^-/-^d2^-/-^ GC^flox/flox^ Cre+ mice, while relatively lower tissue concentrations were associated with polyclonal gammopathy or serum immunoglobulins in the healthy reference range. As determined by serum protein electrophoresis and immunofixation, monoclonal gammopathy was detected in 5 out of 9 CD1^-/-^ GC^flox/flox^ Cre+ mice (Table 1, Fig. 4 B) and was associated with inflammation in lungs, liver and kidneys.

**Table 1.** Monoclonal gammopathy in Gaucher and CD1-/-GCflox/flox Cre+ mice.

| Mice | No treatment |  | Eliglustat |  |
| --- | --- | --- | --- | --- |
|  | M-protein | Total | M-protein | Total |
| CD1<sup>+/+</sup> GC<sup>flox/flox</sup> Cre<sup>+</sup> | 24 | 63 | 0 | 14 |
| CD1<sup>-/-</sup> GC<sup>flox/flox</sup> Cre<sup>+</sup> | 5 | 9 | 3 | 8 |
| CD1<sup>-/-</sup> GC<sup>+/+</sup> | 0 | 8 | - | - |

**Figure 4.**
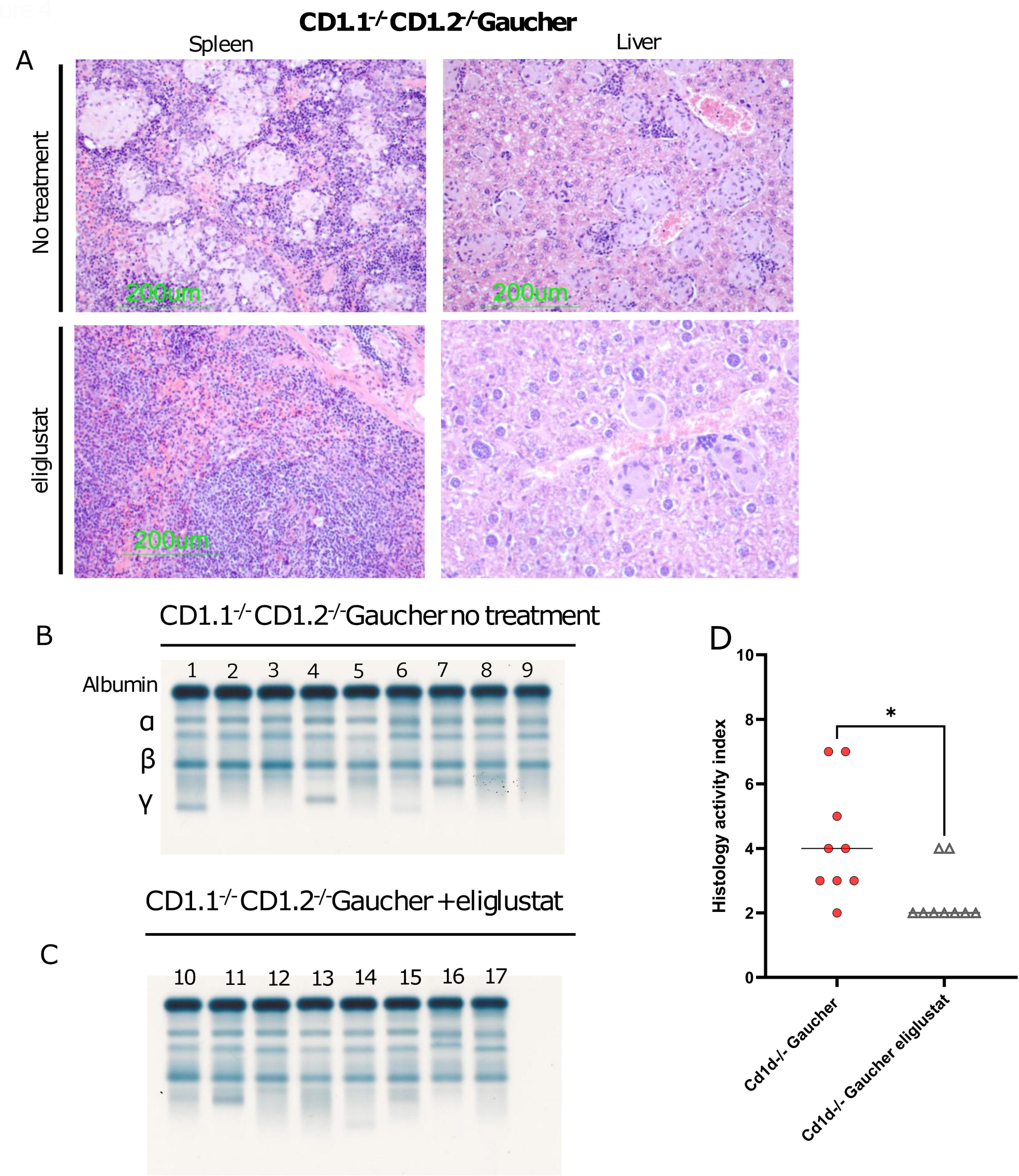
Monoclonal gammopathy and effect of eliglustat on systemic inflammation in CD1d1-/-d2-/-GCflox/flox Cre+ mice. A) Spleen and liver H&E-stained sections of 14 month old untreated and age-matched eliglustat treated mice. Infiltration of enlarged macrophages in spleen and liver parenchyma in untreated mouse. No macrophage infiltration in spleen, reduction of lipid engorged macrophages in liver and less inflammation in eliglustat treated CD1d1-/-d2-/-GCflox/flox Cre+ mice. Protein electrophoresis in serum samples of mice: B) Samples #1, 4, 6, 7, 8 of 14-15 months untreated mice with presence of monoclonal immunoglobulin (n= 9); C) Samples # 16, 17, 21 of age-matched mice treated with eliglustat: monoclonal immunoglobulins are present after treatment. D) Histological activity scores examined in untreated (n=9) and eliglustat treated (n=9) mice. There is a significant reduction of the overall tissue histological activity score in eliglustat treated mice in comparison with age-matched untreated controls (p<0.05). The lines represent Medians, *p<0.05 Mann-Whitney U test.

In control mice from distinct susceptibility strains with identical exposure to the induction stimulus, the background rate of cancer, serological and tissue evidence of autoimmunity was very low. Without inducible Gaucher disease but after exposure to the poly [I:C] inducing protocol, no cancers or autoinflammatory pathology was detectable.

### Absence of autoantibodies in Gaucher mice with CD1d1 and CD1d2 deficiency

Analysis of serum samples did not identify anti-nuclear autoantibodies in 17 CD1^/-^ GC^flox/flox^ Cre+ mice (Table 2, Fig. 3B) and these were not altered after eliglustat exposure.

**Table 2.** Autoantibodies in Gaucher and CD1-/-GCflox/flox Cre+ mice.

| Mice | No treatment |  | eliglustat |  |
| --- | --- | --- | --- | --- |
|  | ANA | Total | ANA | Total |
| CD1<sup>+/+</sup> GC<sup>flox/flox</sup> Cre<sup>+</sup> | 22 | 62 | 3 | 12 |
| CD1<sup>-/-</sup> GC<sup>flox/flox</sup> Cre<sup>+</sup> | 0 | 9 | 0 | 9 |
| CD1<sup>-/-</sup> GC<sup>+/+</sup> | 0 | 8 | - | - |
| CD1<sup>+/+</sup> GC<sup>+/-</sup> | 2 | 9 | - | - |

### Inhibition of glycosphingolipid synthesis in GD mice with CD1d deficiency

To determine if glycosphingolipid-driven immunoglobulin dyscrasia and secretion of autoantibodies reverted upon removal of the antigen, we treated the CD1^-/-^ GC^flox/flox^ Cre+ and CD1^+/+^ GC^flox/flox^ Cre+ mice with eliglustat, a potent and specific orally active inhibitor of UDP-glucose ceramide glucosyltransferase, at the dose of 250mg/kg/day for up to 16 months. No monoclonal paraprotein was detected in CD1^+/+^ GC^flox/flox^ Cre+ (Gaucher) mice after 15 month of eliglustat exposure, thus confirming our previous report of a strong preventative effect of the agent on monoclonal gammopathy associated with Gaucher disease. However, despite significant reduction of *de novo* synthesis of glucosylceramide (Fig. 5, Fig.S1-3), monoclonal gammopathy was detected in three out of eight CD1^-/-^ GC^flox/flox^ Cre + mice (table 1, Fig. 4C) with induced deficiency of glucosylceramidase and accumulation of pathological sphingolipids in visceral organs. To evaluate the effects of long-term (up to 16 months) exposure to eliglustat on development of systemic inflammation and lymphocyte proliferation, tissue sections were evaluated and scored blindly. Hepatic inflammation decreased significantly and no progression to B cell lymphoma/myeloma or other tumours in CD1^-/-^ GC^flox/flox^ Cre+ mice was found. However, 15 GD animals developed hepatitis and cancer (Fig. 4 D, Fig.1F). These data indicate that in addition to the antigenic stimulation, other factors contribute to the adaptive immune response associated with glycosphingolipid accumulation and the outspoken expression of lipid antigens that occur in Gaucher disease. The inflammatory infiltration and polyclonal B-cell proliferation were suppressed in mice receiving eliglustat (Cerdelga®). Of importance retrospective analysis of the archived serum samples of 12 Gaucher mice that had been treated with 300mg/kg of eliglustat in our original study (Pavlova et al., 2015) failed to identify any in which autoantibodies were present.

**Figure 5.**
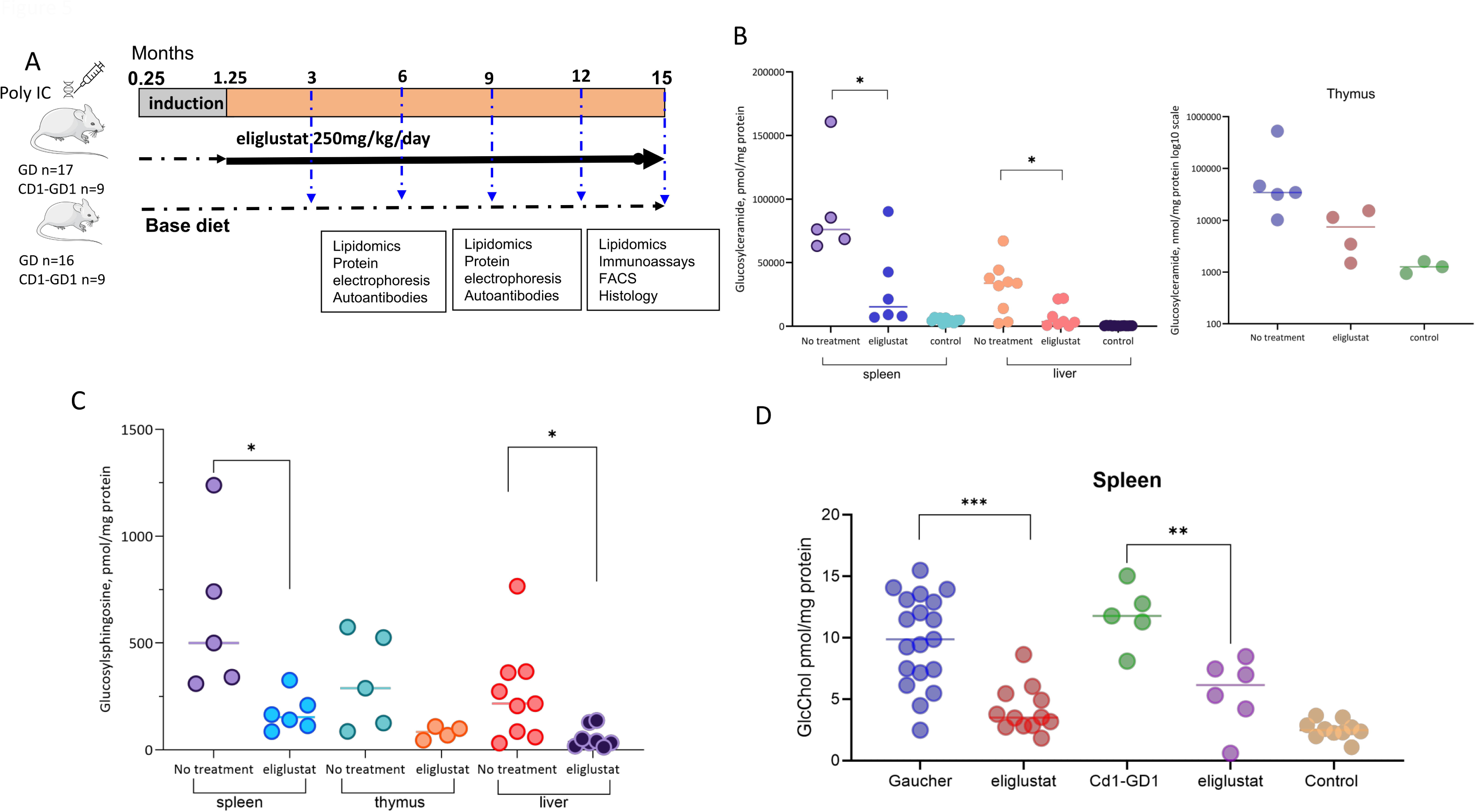
Glycosphingolipid concentrations in Gaucher and CD1d1-/-d2-/-GCflox/flox Cre+ mice. Tissue concentration of glucosylceramide .and glucosylsphingosine in CD1d1-/-d2-/-GCflox/flox Cre+ mice. (A) Schematic illustration of the experimental protocol. B) Glucosylceramide concentration in spleen and liver after treatment with eliglustat was significantly reduced (p <0.05). GlcCer Median concentration in spleen and liver of 15-16 months old untreated mice was 76,194 nmol/mg protein (range 63,191 -160,875) and 33,749 (range 2280 – 67,165) respectively and these were reduced by 60% after 15 months of treatment with eliglustat. GlcCer Median concentration in thymus was 34,630 (range 10,205 – 526,101) and 14,372 (range 1489 – 15,361) after eliglustat exposure. This was not significantly differed compared with age-matched untreated CD1d1-/-d2-/-GCflox/flox Cre+ mice. (C) Glucosylsphingosine concentrations were significantly elevated in all tissues of CD1d1-/-d2-/-GCflox/flox Cre+ and were reduced by 60% in spleens and livers and by 70% in thymus after 12-15 months treatment with eliglustat. D) Increased GluChol concentrations in spleen of GD and CD1d1-/-d2-/-GCflox/flox Cre+. GlcChol in spleen of eliglustat treated mice was reduced by ∼2-fold (<0.05) after 14-15 months of treatment with eliglustat. ns, not significant (p > 0.05); *, p < 0.05. Unpaired two-tailed Student’s t test. Bars and error are mean and SEM.

### Glycosphingolipid Determinations

Tissues and serum β-glucosylceramide and β-glucosylsphingosine concentrations were greatly elevated in Gaucher and Cd1d1^-/-^d2^-/-^ GC^flox/flox^ Cre+ mice: (Fig.5 B and C, Fig. S1 -3). Median concentration in thymus was 34,630 (range 10,205 – 526,101 pmol/mg protein) and 14,372 (range 1489 – 15,361 pmol/mg protein) with eliglustat treatment this was not significantly different in comparison with age-matched untreated Cd1d1^-/-^d2^-/-^ GC^flox/flox^ Cre+ mice. Moreover, glucosylceramide and glucosylsphingosine concentrations were higher in spleen and thymus of Cd1d1^-/-^d2^-/-^ GC^flox/flox^ Cre+ mice with monoclonal gammopathy (range above 200,000 pmol/mg protein). Further to investigate glycosphingolipid metabolism, we assayed hexylceramides and sterylglycosides: HILIC separation of glucosyl and galactosyl-ceramides in tissues of Gaucher and CD1d1-/-d2-/-GCflox/flox Cre+ identified 10% of galactosylceramide out of total HexCer present in the tissues. GlcChol concentration in thymus was increased by 4-fold in spleen or liver (Fig. 5 D, Fig. S1-3).

Analysis of sphingoid bases in thymuses and spleens of Gaucher mice showed decreased long-chain acyl (C24) ceramide species and increased C16-ceramides and dihydroceramides (Fig.6). Total ceramide concentrations were unchanged. Mean sphingosine concentrations were elevated in spleens of Gaucher mice and C16-sphingomyelin and C24-sphingomyelin concentrations were elevated in liver tissue.

**Figure 6.**
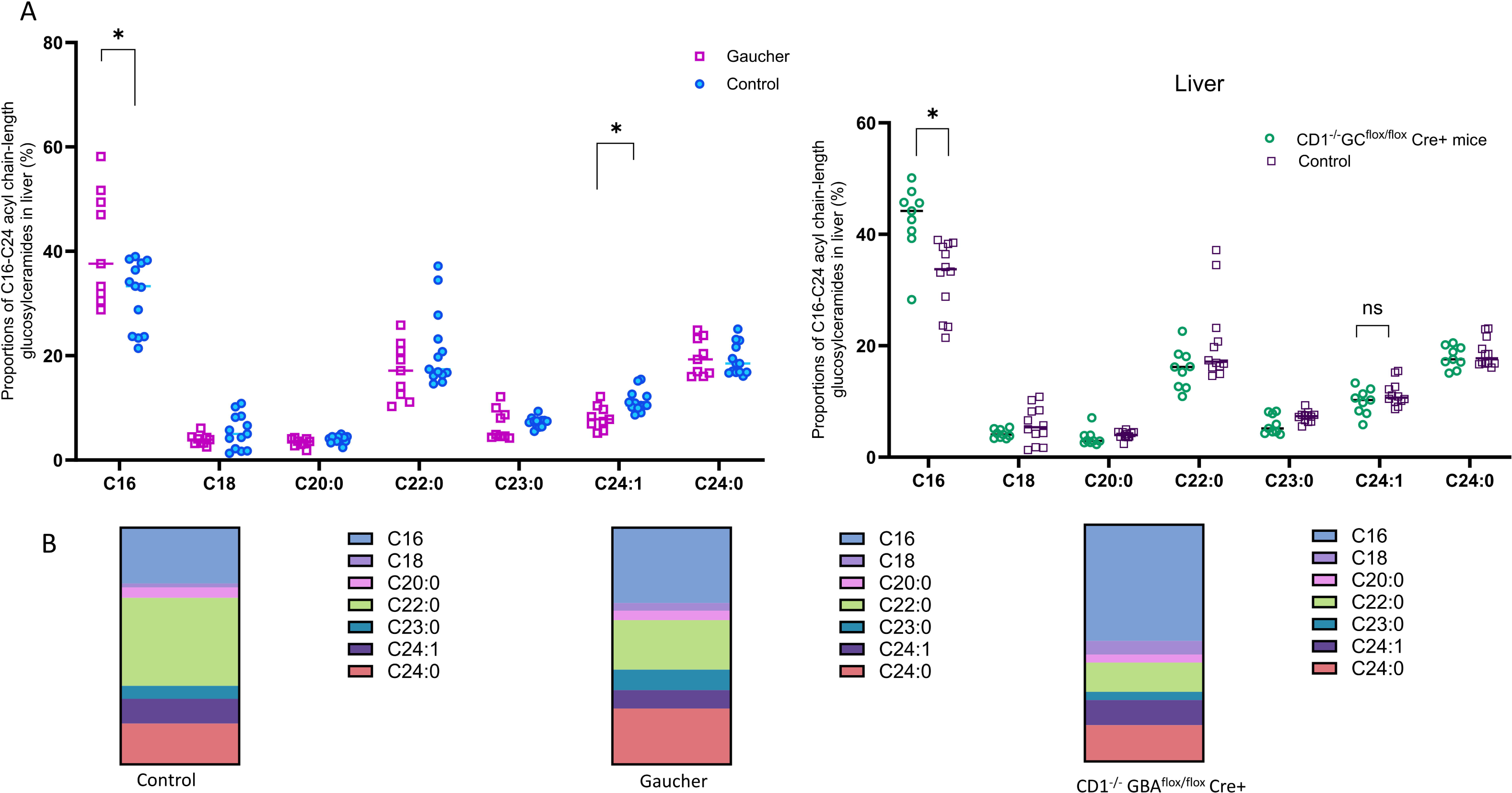
Decreased long-chain acyl (C24) ceramide species and increased C16-ceramides in in liver tissue of Gaucher and CD1d1-/-d2-/-GCflox/flox Cre+ mice. A) Glucosylceramides are presented as proportions of the total GlcCer isoforms per sample based on the length and saturation of their acyl chain. There is decreased C24:1 (long) acyl chain-length ceramides in GD mice (p<0.05), while Cd1d1-/-d2-/-GCflox/flox Cre+ mice have increased C16 acyl chain ceramides in tissues (p<0.05). B) Proportions of specific isoforms in GD, Cd1d1-/-d2-/-GCflox/flox Cre+ and background control mice. ns, not significant, * p<0.05 Mann-Whitney U test. Middle lines are Medians.

### CD1d expression in tissues of Gaucher and CD1d1^-/-^CD1d2^-/-^ GC^flox/flox^ mice

In murine thymus, CD1d is expressed on cortical and medullary thymocytes. The NK1.1+ T cell sub-population in the thymus is comprised of double-positive CD4CD8 thymocytes that undergoes positive selection in the process of NK T cell development. Thymocytes and splenocytes were analysed by flow cytometry to evaluate CD1d expression with the anti-mouse CD1d antibodies (Fig. 3B). CD1 positive cells were detected in control mice and were absent in CD1-/-GCflox/flox mice. Mice heterozygous for CD1 had an intermediate proportion of CD1 positive cells). Somatic mosaic deletion of mutant glucocerebrosidase in sub-population of hematopoietic cells was generated through activation of Cre recombinase with poly [I:C] to induce α-interferon release, as previously described (Pavlova et al., 2013a).

### Autoantibodies in serum of patients with liver Gaucher disease

Polyclonal and monoclonal B cell proliferation with increased serum immunoglobulin concentration is frequent in adults with GD (Nguyen et al., 2020). Although increased autoantibody titres have been previously recognised in patients with Gaucher disease no clear connection has been made with the development of clinical autoimmunity or hepatic cirrhosis (Schoenfield and Mozes, 1990; Serratrice et al., 2018; Lachmann et al., 2000; Regenboog et al., 2018). To investigate if autoantibodies are associated with autoimmune disease as we observed in Gaucher mice, we studied sera from 16 adult patients (13 females and 3 males, mean age 45.1 ± 10.5 years) with clinically confirmed liver cirrhosis or fibrosis collected before or shortly after enzyme replacement therapy was started. We identified an increased frequency of anti-nuclear (ANA), raised anti-ds DNA and antimitochondrial autoantibodies in Gaucher patients with cirrhosis (eight, four and six respectively out of 16 patients) (Fig. 7) suggesting the presence of activated self-reactive cells triggered by pathological lipid metabolism due to lysosomal glucocerebrosidase deficiency. While antinuclear and anti-mitochondrial antibodies can indicate autoimmune hepatitis and cirrhosis, anti-dsDNA IgG antibodies are specific to lupoid autoimmune hepatitis and serve as markers of the severity of the autoimmune state. Moreover, autoantibodies were not frequent in patients with monoclonal gammopathy (Table 3). We retrospectively identified elevated concentrations of all analysed autoantibodies in an archived pre-treatment serum sample of one patient (case 4, in (Lachmann et al., 2000) who developed cirrhosis and hepatocellular decompensation at age of 28 years despite enzyme replacement therapy; this patient had subsequently undergone a successful liver transplantation for which he received immunosuppressive therapy.

**Figure 7.**
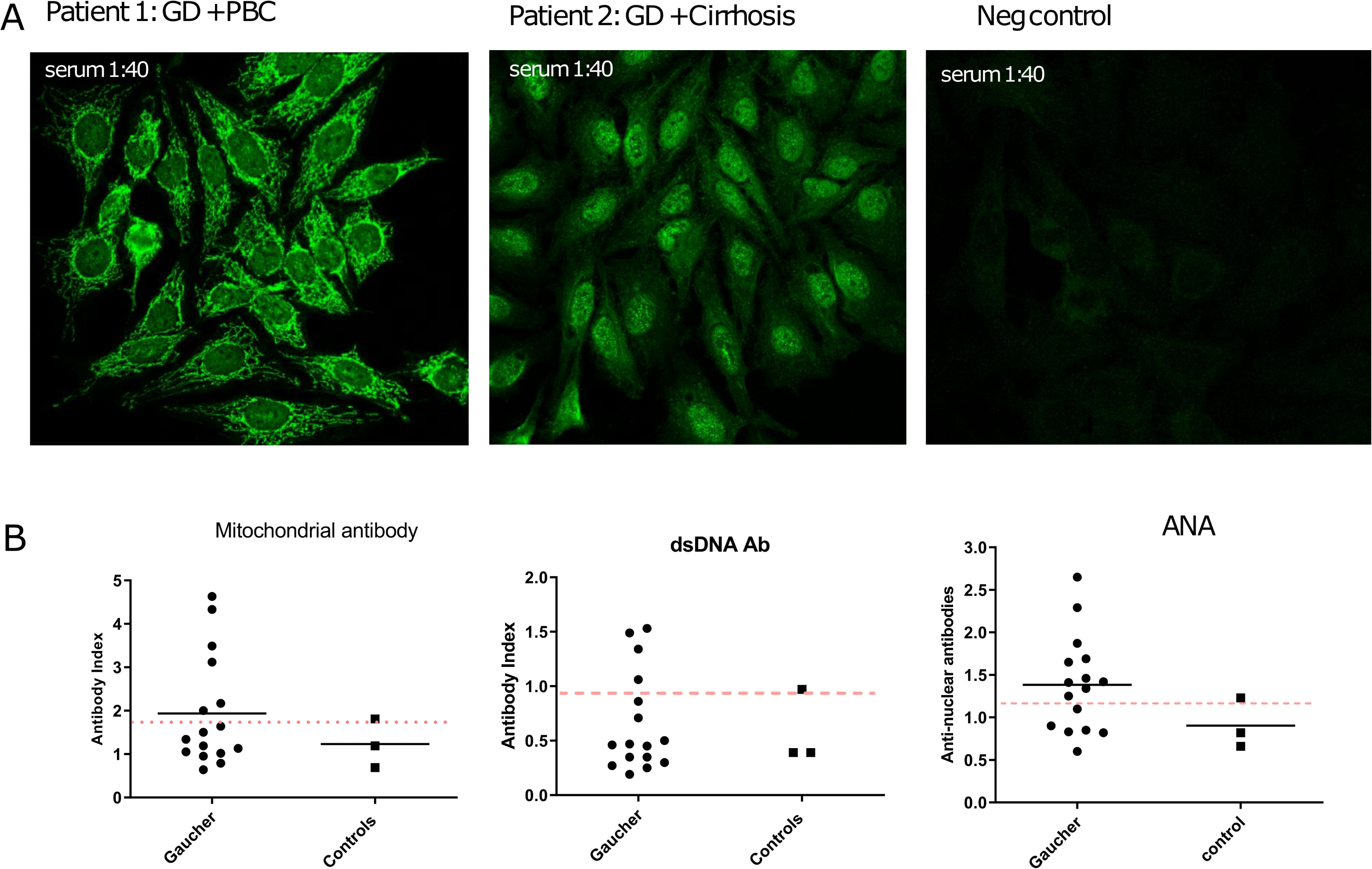
Prevalence of autoantibodies in patients with cirrhosis and Gaucher disease. A) Representative autoantibodies patterns observed by indirect immunofluorescence on HEp-2 cells identified in serum of patients with Gaucher disease and cirrhosis. Sera were used to stain Hep-2 cells, and the presence of autoantibodies was detected by anti-human Ig G conjugated to FITC and confocal microscopy. The bright nuclear staining of Hep-2 cells indicates that patient #1 sera bind to nuclear antigens. Sera of patient 2 are immunoreactive to mitochondrial antibodies evidenced by cytoplasmic pattern to primary biliary cirrhosis. B) Mitochondrial, dsDNA and anti-nuclear antibodies in patients with Gaucher disease with cirrhosis and healthy controls. The lines indicate medians. The pink dotted lines show normal range.

**Table 3.** Relationship between presence of autoantibodies, monoclonal gammopathy and hepatic cirrhosis in patients with Gaucher disease.

| Gaucher disease |  | N autoantibodies-<br>positive | N paraprotein<br>positive | Mean, age<br>(years) |
| --- | --- | --- | --- | --- |
| Cirrhosis | 11 | 7 | 0 | 44.9 ±10.8 |
| Liver fibrosis | 5 | 1<sup>†</sup> | 1 | 45.6 ± 10.9 |
| Total | 16 | 8 | 1 |  |

<sup>†</sup> -anti-ds DNA antibody positive

In the Gaucherite cohort, 17 out of 201 patients had monoclonal gammopathy, 15 had positive anti-nuclear antibodies, 4 of these patients with ANA had monoclonal gammopathy and in two, anti-DNA antibodies were found.

To further investigate if autoantibodies in patients’ sera target liver-specific antigens, clinical immunoblot analysis was carried out in a set of pre-selected serum samples. Most samples were immunoreactive to the selenocysteinyl-tRNA(Sec) synthase, classified as a soluble liver antigen/liver pancreas (SLA/LP), which is highly specific for type I autoimmune hepatitis; in addition the autoantibodies were reactive to the core structure of the pyruvate dehydrogenase complex including M2-3E (BPO), PDH M2, the nuclear membrane proteins GP-210 and SP100 which are exclusively found in primary biliary cirrhosis (Table 4).

**Table 4.** Distribution of liver specific autoantibodies in patients with Gaucher disease.

| Liver-related autoantibodies | N autoantibodies-<br>positive |
| --- | --- |
| PDH (M2) | 1 |
| M2-3E(BPO) | 1 |
| SP100 (ANA) | 1 |
| PML (ANA) | 0 |
| GP210 (ANA) | 2 |
| Liver kidney microsomal-1 (LKM-1) | 0 |
| Liver cytosolic antigen 1 | 1 |
| Soluble liver antigen/liver pancreas | 6 |
| Anti-ds DNA antibody | 2 |
| Crithidia luciliae ds DNA | 1 |

## Discussion

Our investigations in murine models of Gaucher disease provide insights into a sustained immune reaction that is triggered by excess glycosphingolipids. We contend that the findings are relevant to the pathogenesis of human Gaucher disease and emphasise the importance of glycosphingolipids in the development of autoimmunity and cancer. In addition to monoclonal gammopathy, we report a severe autoimmune hepatitis and autoinflammation in experimental mice with a chimeric F1 mixed background. Unexpectedly, the prominent liver inflammation occurred in a strain of these mice with inducible Gaucher disease: there were pathognomonic histological and immunological features of chronic active hepatitis. While the molecular pathways that drive autoimmunity and monoclonal gammopathy in Gaucher disease are ill-understood, development of B cell proliferation with monoclonal gammopathy, B cell cancers, liver disease and liver cancer with autoimmunity in the murine models studied here are conditional upon induction of Gaucher disease and the pathognomonic hallmarks of defective glucosylceramide recycling.

Affected animals have greatly elevated β-glucosylsphingosine and β-glucosylceramide concentrations in several tissues. We further explored the metabolic fate of these glycosphingolipids by examining the proposed transglucosylation of glucosylceramide on the plasma membrane and cytoplasm, catalysed by the neutral glucosylceramidase, GBA2, which generates glucosylcholesterol (GlcChol) (Marques et al., 2016); Boer et al., 2021). Splenic tissue from Gaucher mice and Gaucher mice lacking CD1 had markedly elevated GlcChol and the GlcChol concentration was 4-times greater in thymic tissue, and may be associated with an enhanced metabolic rate and apoptosis in this organ. Moreover, further analysis of the ceramide isoforms identified reduced C24:1 and C22:0 acyl chain-length ceramides and increased C16 acyl chain ceramides in serum and tissues. While the membrane trafficking and metabolic degradation of these lipids are not well characterized, sphingolipid analysis shows that in mice with GBA1 deficiency, greatly increased β-glucosylsphingosine derived from glucosylceramide and the elevated β-glucosylceramides are a mixed population derived from *de novo* synthesis and resynthesis through the ceramide salvage pathway.

Impaired lysosomal recycling of glucosylceramide and reduced synthesis of long-chain C24-ceramides by the salvage pathway appears to induce compensatory activation of de novo C16-ceramide biosynthesis by CerS2 and/or CerS6 (Park et al., 2014).

Systemic injury, including exaggerated proliferation in the liver, results from sustained autoimmune attack: we contend that the altered sphingolipid milieu of Gaucher disease impedes access of immunoregulatory T cells and disrupts control of tolerance in the periphery. Membrane lipid biology and trafficking, associated with self-reactive B cell proliferation and autoimmunity merit further study to understand the roles of the lipid degradation in immune cell differentiation as well as lipid antigen presentation.

The family of CD1 proteins are a group of cell membrane-associated molecules at the cell surface that partially resemble MHC class I proteins: their α1, α2 and α3 transmembrane domains interact with β2-microglobulin but they present diverse lipids, glycolipids and hydrophobic peptides to T cells for immune surveillance. Based on sequence homology, CD1 molecules are classified to three groups: human group I comprises CD1a, -b and -c; group II, consists of CD1d, and group III, only CD1e. However, different mammals have distinct CD1 isoforms and mice express only CD1d1 and CD1d2 proteins. Unlike the highly polymorphic MHC molecules, structural variation is restricted in CD1 molecules but all bind lipid-based antigens that enable selection and differentiation of Natural killer T (NKT) cells in the thymus and periphery.

These unconventional T cells also express natural killer cell receptors: they are involved in innate immunity but their lineage commitment and differentiation pathways are a subject of debate (Georgiev et al., 2016). Two NKT cell types are defined by the conserved molecular arrangements of their cognate T cell antigen receptor (TCR) α-subunit genes that differ between mammals. The most studied (invariant) iNKT cells recognise the canonical experimental ligand, α-galactosylceramide. Antigenic stimulation leads to release of Th1, Th2, and Th17, IL10 cytokines, and stimulation of cytotoxic responses. Not only are Type I NKT cells activated by bacterial ligands, they sense host changes in endogenous lipids. Potentially relevant to the glycosphingolipid disorder Gaucher disease, is the sustained stimulation of iNKT cells by autochthonous glycolipid antigens that stimulate the release of cytokines which influence the development and chronic activation of vicinal immune cells. CD1d expressed on the surface of antigen-presenting cells can present the endogenous lipid self-antigens – β-D-glucosylceramide and β-glucosylsphingosine, which are selectively over-abundant in Gaucher disease, for recognition by T cell receptors. This would drive an antibody-mediated immune response (Nair et al., 2015b, 2018) and has been presented as a source of antigen-driven myeloma in Gaucher disease.

In later studies, glycosphingolipid binding was demonstrated in single tumour cells isolated from Gaucher patients with gammopathy and in patients previously found to have the sporadic lipid-reactive disorder. The clonal nature of the immunoglobulins was confirmed by protein-sequencing. In patients with lipid-reactivity, gene-expression profiling and cytogenetic studies showed an enriched population of nonhyperdiploid tumours that secreted glycolipid-reactive monoclonal antibodies on exposure to Gaucher-related lipids (Nair et al., 2018).

In our study, the loss of CD1d function in the autoimmune Gaucher mice led to reduction of systemic inflammation but lupus like signs appeared with abundance of storage multinucleated macrophages (Gaucher cells). Cytokines, secreted by these cells contribute to the immune response directed to inflamed or malignant cells in the pathologically increased glycosphingolipid milieu. A further notable observation in the poly [I:C]-(α-interferon) inducible Gaucher mice with CD1d deficiency indicated the evolution of an adaptive immune response to lymphoma in the 129/sv x B6 x BALB/c strain with progressing allogenic recognition. When CD1d1 was deleted in autoimmune-prone BALB/c mice, anti-DNA and anti-phospholipid antibody titres were reduced along with improvement of the lupus however, as in our experiments, the monoclonal immunoglobulins persisted, suggesting a dual regulatory role of the receptor in autoimmunity.

In the animals reported here, eliglustat decreased the signs of hepatic inflammation and lymphoproliferation and monoclonal gammopathy confirming our previous observation and the preventative effect of this inhibitor now in widespread clinical use (Pavlova et al., 2015). By correcting glycosphingolipid-driven autophagy of osteoclastic differentiation factor TRAF3, eliglustat improves osteolytic bone loss in preclinical models of myeloma (Leng et al., 2024, 2022; Ersek et al., 2015) and the less specific nonclinical inhibitor, P4 related to eliglustat, was reported to be cytostatic and cytotoxic to murine myeloma cells (Manning and Radin, 1999).

Invariant natural killer T cells are abundant in the liver and participate in immune surveillance of external pathogens and autologous -malignant cells that migrate through endothelial barriers. CD1d is expressed on endothelial cells of the sinusoids and in the liver resident macrophages, Kupffer cells. Presentation of pathological self-derived sphingolipids by these cells may activate hepatic iNK T cells resulting in autoimmunity (Mattner et al., 2008). Tissue residential iNK T cells and macrophages are enriched in the liver and spleen, which are grossly enlarged in Gaucher disease. However, the liver enlargement is usually accompanied by modest elevation of biliary tract enzymes and liver-related transaminase activities released into the serum (James et al., 1981). While extensive fibrosis progressing to cirrhosis that may require liver transplantation may occur in Gaucher disease, (Ida et al., 1996; James et al., 1981; Lachmann et al., 2000; Nascimbeni et al., 2018), the relationship between pathological accumulation of glycosphingolipids and extensive destruction of liver architecture with infiltration by activated macrophages in severe cases is ill-understood.

To determine whether the autoimmune phenomena observed in our experimental mice occurred in human Gaucher disease, we examined sera obtained from patients in the pre- or early treatment period and those with evidence of hepatic cirrhosis: an unexpectedly high frequency of tissue-specific autoantibodies was found. We therefore conclude that in patients with severe hepatic involvement associated with Gaucher disease including primary liver cancers and cirrhosis, may also be the result of autoimmunity. We contend that as in the mice, pathologic glycosphingolipid abnormalities due to GBA deficiency induce loss of immune tolerance and activate tissue-specific autoimmunity.

Autoantibodies specifically associated with liver disease in patients with Gaucher disease have been reported very rarely (Patel et al., 1986). This may reflect the generally recognised polyclonal increase in immunoglobulins and overall the presence of elevated titres of organ-specific autoantibodies in patients with Gaucher disease, but in several studies, compared with matched control subjects, no clear link to clinical autoimmune disorders has been established (Serratrice et al., 2018; Shoenfeld et al., 1995).

As to environmental factors and autoimmune phenomena, bacteria in the genus *Sphingomonas* are unusual it that they express cell wall glycosphingolipids instead of lipopolysaccharide, typical of other gram-negative bacteria: the glycosphingolipids of the waterborne *Novosphingobiobium aromatacivorum* appear to act as a molecular mimetic that triggers autoimmunity in genetically predisposed patients who develop a phenocopy of primary biliary cirrhosis (Kaplan et al., 2004). Glycosylceramides present in the cell wall of *Sphingomonas sp*. activate host NKT cells via interactions with CD1 molecules and on acute exposure induce a condition analogous to endotoxin shock with spectacular release of ɤ-interferon and largely independent of Toll-like receptor recognition or the presence of LPS. In connection to the prevalent human autoimmune disease, primary biliary cirrhosis the diagnostic hallmark of antimitochondrial antibodies, the phylogenetically related *Novosphingobium sp.* express elements of the conserved mitochondrial pyruvate dehydrogenase complex elements that are highly homologous to the human PDC-E2 (Mattner et al., 2008). These authors identified molecular mimicry in these glycolipids that bind to CD1d glycoproteins in the lysosomal compartment and are recognized by conserved human (and murine) T cell receptors thereby serving as an innate pathway for detecting foreign cell wall components and microbial clearance after infection. NKT cells home to the liver and apparently patrol sinusoidal endothelial cells. Not only are they increased in the liver of patients with primary biliary cirrhosis but increased hepatic CD1d expression is a feature of this disease. Kaplan et al., (2008) showed that mice harbouring *N. aromaticivorans* developed antimitochondrial IgG antibodies and chronic bile duct inflammation in a CD1d-dependent manner accompanied by lymphoepithelioid granulomas that resemble primary biliary cirrhosis. Once established the disease proved to be transferable to naive mice by T cells, independently of CD1d, NKT cells, or microbial infection, thereby establishing autonomous autoimmune disorder.

Previous attempts to immunize autoimmune-prone BALB/c mice with purified anti-DNA antibodies extracted from serum of GD patients failed to induce experimental systemic lupus erythematosus. This led to the conclusion that the high concentrations of autoantibodies present in Gaucher sera represent non-pathogenic polyspecific autoantibodies and are apparently unrelated to autoimmune disease (Shoenfeld and Mozes, 1990; Serratrice et al., 2018).

Here we describe the first experimental model of CD1d1/Cd1d2 deficiency in a conditional inducible murine model of Gaucher disease with *GBA1* deficiency in hematopoietic cells. Overall, our study in Gaucher mice crossed with CD1d-/-mice on an autoimmune background, points to an immune regulatory role of the CD1 in the recognition of the pathologic glycosphingolipids in Gaucher disease. Neither lymphomas nor plasmacytomas were found in animals with CD1 deficiency or those that had received eliglustat that specifically depleted the pathologically over-abundant glucosylceramides and their congeners. The role of CD1 in the generation of CD1-restricted T cell responses requires non-covalent association with β-2 microglobulin on the surface of immune cell membranes where they encounter exogenous lipid antigens and are subject to cross-linking. Murine CD1 molecules traffic to endosomal and lysosomal compartments where they encounter endogenous glycolipids - in the case of β-glucosylceramide, the lipids are loaded in the presence of membrane-associated saposin C that is required for enzymatic cleavage of naturally occurring glucosylceramide by GBA (Darmoise et al., 2010).

We therefore contend that the GBA1 deficiency is associated with severe liver disease and hepatic cirrhosis - a rare but life-threatening complication of human Gaucher disease that appears to account for the increased risk of hepatocellular carcinoma (Regenboog et al., 2018). CD1d deficiency in Gaucher mice reduced the appearance of autoantibodies but monoclonal immunoglobulins were present. The accumulated lipids drive chronic inflammation, autoantibody production, and an increased susceptibility to conditions like B-cell lymphoma and myeloma. Furthermore, eliglustat may be able to mitigate the risk of such comorbidities in patients with early manifestations or disease, as previously reported in murine models and here studied in different informative strains.

## Methods

### Gaucher and Cd1d mice

Mice with inducible mosaic expression of beta-glucocerebrosidase deficiency in haematopoietic cells: excision of exons 9-11 of the murine GBA1 gene [129S1/Sv * 129X1/SvJ * C57BL/6 * CBA Gba tm1Karl/tm1Karl Tg(Mx1-cre)1Cgn/0 (MGI:3687965; MGI:2176073)] were bred by heterozygous mating (Enquist et al., 2006). After multiple generations, the mice were crossed with congenic C57BL/6 mice. All offspring received five peritoneal injections of the interferon inducer polyinosinic - polycytidylic acid [poly(I:C)] acid sodium salt (cat. no. P1530, Sigma) which was dissolved in sterile phosphate-buffered saline (PBS) at concentration of 1 mg/ml, incubated at 50^°^C and cooled at room temperature to ensure re-annealing of double-stranded poly (I:C) molecules; 50–250 µl (10 ug/g mouse weight) of this solution was administered to neonatal mice after 7 days of age. Genotyping was determined as previously described (Pavlova et al., 2013a). Strain control mice GC flox/wt and wt were induced with polyinosinic-polycytidylic acid [poly(I:C)] and maintained for 24 months.

Mice homozygous for the Cd1tm1Gru targeted mutation are deficient in both the Cd1d1 and Cd1d2 genes (CD1.1, CD1.2) (Smiley et al., 1997). Homozygous mutant mice lack the normal natural killer cell -like T cell subset. This strain originated on a 129S background and has been backcrossed to BALB/cJ (N11). It is maintained by homozygous sibling matings. Two breeding pairs, Jax Lab Stock number 003814 B6.129S2-Cd1^tm1Gru^ (Homozygous for Cd1tm1Gru x Homozygous for Cd1tm1Gru) were obtained. Both the Cd1d1 gene and the adjacent Cd1d2 gene were mutated in this allele. The coding sequence of Cd1d1 was replaced with a neomycin selection cassette and the coding sequence of Cd1d2 was deleted; however, the intergenic sequence was retained. Flow cytometric analysis on thymocytes derived from homozygous mice confirmed that no stable encoded protein was expressed on the cell surface (Smiley et al., 1997). Genotyping was carried out in our laboratory and using a real time PCR assay with specific probes designed for each gene in the Transnetyx fully automated system (Transnetyx, Cordova, TN).

The Mouse Universal Genotyping Array (MUGA), MiniMUGA, an array-based genetic QC platform with over 11,000 probes was used to determine the genetic background of the mouse strains used (Sigmon et al., 2020).

### Substrate reduction therapy with eliglustat

Gaucher mice received eliglustat tartrate admixed in a standard rodent LabDiet Rodent Diet, (TestDiet, USA). A group of 17 Gaucher and 9 Cd1d1-/-d2-/-GCflox/flox Cre+ mice received 250 mg/kg daily, and 16 Gaucher and 9 Cd1d1-/-d2-/-GCflox/flox Cre+ mice received standard diet LabDiet Rodent Diet (TestDiet). Eliglustat was administered *ad libitum* from 6 weeks (1.5 months) of age for up to 16 months. After two weeks of exposure activity of the drug was measured by glucosylceramide and glucosylsphingosine concentration in serum samples of treated mice. Health and body weight monitoring performed regularly. A humane end point was defined as the loss of 15% of greatest achieved total body weight or signs of sickness. The animals were housed in individually ventilated cages in groups of up to four mice. Blood samples were collected from the lateral saphenous vein at baseline and in three-months intervals. Serum samples were stored at -70 until analysis.

Body Condition Scoring was used for assessing endpoints: abdominal swelling and muscle loss (amended from Office of Ethics and Compliance Institutional Animal Care and Use Program, IACUC Standard Procedure) (Ullman-Culleré and Foltz, 1999; IACUC Guidelines, University of California at San Francisco).

The regulated experimental procedures were approved by the UK Home Office under the Animals Scientific Procedures Act (1986) licence number P09AD3ABE.

DNA from ear biopsy sample and from tissues homogenates was isolated using DNeasy Blood & Tissue Kit (catalogue # 69504, Qiagen) and used for genotyping and genetic analysis. RNA from spleen and liver homogenates was purified using RNeasy Mini Kit (Qiagen) following the manufacturer’s guidance.

### Histology and immunohistology

Spleen, liver, thymus, lungs and kidneys were collected by dissection and fixed in 4% w/v paraformaldehyde and processed in paraffin-embedded blocks. Paraffin-embedded sections were stained with haematoxylin and eosin (H&E) or Masson Trichrome. Immunohistochemical staining was performed using anti-mouse CD45R/B220 (R and D Systems Cat# MAB1217, RRID:AB_357537), CD3Ɛ (Abcam Cat# ab16669, RRID:AB_443425), CD138 (BD Biosciences Cat# 553712, RRID:AB_394998), Ki-67 (Abcam Cat# ab15580, RRID:AB_443209) antibodies.

Biotin-SP-conjugated donkey anti-rat IgG (Jackson ImmunoResearch Labs Cat# 712-066-153, RRID:AB_2340649) and anti-rabbit IgG (Jackson ImmunoResearch Labs Cat# 711-066-152, RRID:AB_2340594) were used as secondary antibodies. Immunostaining was developed using avidin-biotin peroxidase and diaminobenzidine technique (Vectastatin ABC HP Kit, Vector Laboratories) and counterstained with haematoxylin.

### Flow cytometry analysis

Thymuses, spleens, lymphomas were collected from untreated and eliglustat-treated Gaucher mice at 14-15 months of age. Single cells suspensions were prepared by mechanical dissociation using a 70 µm cell strainer (BD Falcon), cells were stained on the same day of collection or resuspended in cryopreservation media and stored in liquid nitrogen until analysis. Erythrocytes were lysed in spleenocytes suspensions using Lysing Buffer for 10 minutes in RT. Dead cells were discriminated by staining with Zombie UV fixable viability dye (Biolegend) in PBS, according to manufacturer’s instructions and were excluded from the downstream analysis. Fc block was used to prevent non-specific binding to Fcγ receptors. Single cell suspensions were stained with the following antibodies: anti-mouse antibodies CD45R/B220 (Thermo Fisher Scientific Cat# 15-0451-82, RRID:AB_468752), CD1d (Thermo Fisher Scientific Cat# 64-0011-82, RRID:AB_2688078), NK1.1 (Thermo Fisher Scientific Cat# 64-5941-82, RRID:AB_2662737), CD4 (BD Biosciences Cat# 563727, RRID:AB_2728707), CD8α (130-097-025, Miltenyi Biotec), TCR V β 10 ((Thermo Fisher Scientific Cat# 11-5805-80, RRID:AB_2043882), CD25 (BD Biosciences Cat# 561257, RRID:AB_10611871), CD44 (BioLegend Cat# 103027, RRID:AB_830784), CD69 (Miltenyi Biotec Cat# 130-103-946, RRID:AB_2659083), IgM (BD Biosciences Cat# 563837, RRID:AB_2869524), IgD (BD Biosciences Cat# 565348, RRID:AB_2739201), CD23, CD21, CD19, CD3, CD11b. Analysis was performed using BD Fortessa Analyser (BD Biosciences) and analysed with FlowJo software (BD Biosciences

### Autoantibody profile in mice

Serum from GD mice were used to stain human endothelial cells (HEP-2) that used as the antigenic substrate (MBL Bion and BioRad) in indirect fluorescent antibody assay. S. Patterns of anti-nuclear autoantibodies identified by indirect immunofluorescence on Hep-2 cells stained with serum from Gaucher or CD1d-/-GCflox.flox Cre+ mice. For initial screening serum dilution was 1:40. Serial dilution up to 1:360 with PBS was carried out to determine autoantibodies titres in a subset of samples. Negative and positive staining controls were used in each analysis as an internal reference control. Slides were stained with anti-mouse IgG-Alexa 488 and visualised on a Leica SPE confocal microscope.

Anti -ss-DNA, ds-DNA, smooth muscle, anti-mitochondrial autoantibodies were determined in serum samples of mice using commercial ELISA kits.

### Glycosphingolipid concentrations

Lipidomicanalysis was carried out in three independent laboratories:

1. <u>Lipidomics Shared Resource, Hollings Cancer Center, Medical University of South</u> <u>Carolina.</u> Sphingolipids were extracted by the previously described methods.Ceramidesand other sphingolipids were measured by high-performance liquid chromatography mass spectrometry (LC-MS/MS) methodology as previously described Concentrations of glycosphingolipids were normalised to total protein concentration.
2. Medical Biochemistry Department, Leiden University, The Netherlands. GluCer, GluSph, GalCer, GalSph, GlcChol and GalChol concentrations were quantified in serum and liver, spleen and thymus tissues lysates by mass spectrometry as previously described ( Lelieveld et al., 2019) Neutral (glyco)sphingolipids, (glyco)sphingoid bases, and hexosylcholesterol (HexChol) were extracted using an acidic Bligh and Dyer procedure [1:1:0.9 chloroform:methanol:100 mM formate buffer (pH 3.1)] according to methods described before (A, B, C). To a sample was added 20 μl of internal standard mixture (0.1 pmol/μl of 13C5-sphinganine, 13C5-sphingosine, 13C5-GlcSph, 13C5-lysoGb3, and C17-lysoSM in methanol), 20 μl of C17-dihydroceramide (20 pmol/μl in methanol), 20 μl of 13C6-GlcChol (0.1 pmol/μl in methanol) followed by methanol and chloroform (2:1, v/v). After brief mixing, the samples were left at room temperature for 1 h with occasional stirring and 3× 1 min sonication in a bath sonifier (VWR ultrasonic cleaner USC). Samples were centrifuged for 10 min at 15,700 g to spin down precipitated proteins. The supernatant was transferred to a clean tube, while excess organic solvent was evaporated. Chloroform and 100 mM formate buffer (pH 3.1) were added to the supernatant, to a final ratio of 1:1:0.9 methanol:chloroform:formate buffer, to induce separation of phases. The upper phase was used for analysis of lyso(glyco)sphingolipids and the lower phase for analysis of neutral (glyco)sphingolipids and HexChol. After centrifugation, the upper phase was transferred to a clean tube and the lower phase (chloroform phase) was extracted an additional time with methanol and formate buffer. Pooled upper phases were concentrated at 45°C in an Eppendorf Concentrator Plus and a butanol/water (1:1, v/v) extraction was performed. The upper phase (butanol phase) was transferred to a clean tube and concentrated. Lipids were dissolved in 100 μl of methanol, stirred, sonicated for 30 s in a bath sonicator and centrifuged. The supernatant was transferred to a vial for subsequent LC-MS/MS analysis. The remaining lower chloroform phase was transferred to a clean tube and the interphase was washed with chloroform. The pooled lower chloroform phases were split, whereby one part was used to analyze HexChol and the part for analysis of neutral glycosphingolipids was transferred to a Pyrex tube and dried at 45°C under a gentle stream of nitrogen. Deacylation was performed by adding of 500 μl of sodium hydroxide (0.1 M NaOH in methanol) using a microwave-assisted saponification method (50). The samples were cooled and neutralized by adding hydrogen chloride (50 μl of 1 M HCl in methanol) and dried, followed by butanol/water extraction and preparation for LC-MS/MS as described above. For determination of HexChol (A), the other half was concentrated; a butanol/water extraction was performed; and samples were prepared for LC-MS/MS analysis as described above. For hydrophilic interaction liquid chromatography (HILIC) separation lipids were extracted as described above and lipids were resuspended in acetonitrile:methanol (9:1, v/v) prior to transfer to LC-MS vials [D]. A BEH HILIC column (2.1 × 100 mm with 1.7 μm particle size; Waters Corporation) was used at 30°C for the separation of lipids with glucosyl and galactosyl moiety. In general, eluent A contained 10 mM ammonium formate in acetonitrile/water (97:3, v/v) and 0.01% (v/v) formic acid, and eluent B consisted of 10 mM ammonium formate in acetonitrile/water (75:15, v/v) and 0.01% (v/v) formic acid. Lyso- and deacylated glycosphingolipids were eluted in 18 min with a flow of 0.4 ml/min using the following program: 85% A from 0 to 2 min, 85–70% A from 2 to 2.5 min, 70% A from 2.5 to 5.5 min, 70–60% A from 5.5 to 6 min, 60% A from 6 to 8 min, 60–0% A from 8 to 8.5 min, 0–85% A from 8.5 to 9.5 min, and re-equilibration of the column with 85% A from 10 to 18 min. HexChol was eluted in 18 min with a flow of 0.25 ml/min using the following program: 100% A from 0 to 3 min, 100–0% A from 3 to 3.5 min, 0% A from 3.5 to 4.5 min, 0–100% A from 4.5 to 5 min, and re-equilibration with 100% A from 5 to 18 min. Data were analyzed with MassLynx 4.1 software (Waters Corporation).
3. Rare and Neurologic Disease Research, Sanofi R&D. Tissue samples were prepared as reported previously (Zaccariotto et al., 2022)(Zaccariotto et al., 2022) (Eva Zaccariotto et al. Biomedicine & Pharmacotherapy 149 (2022) 112808). Quantitative analysis of sphingolipids was performed by liquid chromatography and tandem mass spectrometry (LC/MS/MS). Briefly, an aliquot of 10 μl of serum or tissue homogenate was extracted with 1 ml of 50:50 acetonitrile/methanol (v/v) containing internal standards (C12-galactosylceramide at 5 ng/mL or D5-labeled glucosylsphingosine at 1 ng/mL). Extract from each sample was transferred into a 384-well microtiter plate for analysis. Glucosylceramide and galactosylceramide were separated by a Cortecs HILIC column (2.1×100 mm, 2.7 µm particle size, Waters Corp., Milford, MA) and analyzed using a Vanquish UHPLC (Thermo Fisher Scientific, West Palm Beach, FL) coupled to an AB Sciex API 5000 triple quadrupole mass spectrometer (Applied Bio systems, Foster City, CA). For MRM transitions, Q3 was set to detect ions with m/z 264.2, while Q1 was set to detect ions with m/z 644.5, 700.6, 728.6, 756.6, 784.6, 798.6, 810.8, and 812.8. Glucosylsphingosine and psychosine were separated by a Waters BEH HILIC column (2.1 mm × 100 mm, 1.7 m particle size) and analyzed using a Vanquish UHPLC coupled to an AB Sciex API 6500 Plus triple quadrupole mass spectrometer. For the MRM transition, Q1 and Q3 were set to 462.3 and 282.3, respectively. The calibration curves were constructed using C16:0 glucosylceramide or glucosylsphingosine standards (Avanti polar lipids, Inc., Alabaster, Al).

### Serum protein electrophoresis

Serum protein electrophoresis was performed in agarose gels using semi-automatic gel electrophoresis instrument (Hydrasys, Sebia) in 10µL samples of serum. Samples with abnormal electrophoretic patterns were analysed to electrophoresis with immunofixation using anti-mouse antibodies against heavy and light immunoglobulin chains. The gels were subjected to densitometry analysis using optical gel scanner (Sebia). Total serum protein concentration was determined by colorimetric assay based on a bicinchoninic acid procedure (Protein Assay, Thermo Scientific).

#### Total serum immunoglobulin concentrations

Total Ig G, M ELISA kits (Affymetrics, UK) were used to analyse serum immunoglobulins in groups of GD and control mice.

### Gaucher disease patients and controls

A cohort of 16 patients Gaucher disease who attended the Cambridge University Hospital were studied. Venous blood was obtained from each patient at the time of outpatient clinic attendance. Serum was transferred to polypropylene tubes and frozen at −80 °C until required. Retrospective non-identifiable clinical data were analysed. Informed consent was obtained from the parents of the patients in accordance with the standards of the Declaration of Helsinki. This aspect of the study was approved by the Ethics Committee of University of Cambridge. In addition, we analysed serum samples of 201 GD patients recruited in the UK MRC Gaucher Investigative Therapy Evaluation (Gaucherite) Consortium. Ethical approval was given by the national ethics review committee (14/EE/1168) under the purview of the NHS Research and Development (R&D) in the United Kingdom; sera were stored in the biobank curated in the Department of Pharmacology, University of Oxford (Prof F Platt and Dr Kerri Wallom). The establishment, and clinical characteristics of the patients have been reported elsewhere; the database is hosted by the University of Cambridge (D’Amore S et al, 2021).

### Autoantibody analysis in human serum samples

Immunological assays for autoantibodies were carried out in our laboratory and by clinical diagnostic services at UKAS certified laboratories using commercially available liver Autoantibodies HEP-2 ANA, Anti-Mitochondrial (AMA), Smooth muscle antibodies (SMA), Anti-Gastric Parietal cell antibodies (GPC), Liver kidney microsomal antibodies (LKM)(LAB2165). Anti DNA LAB2018 (DNA(ds)) antibody ELISA). Specific liver autoantibodies [LAB 2165] and anti-double stranded DNA antibody ELISA were carried out. Sera were obtained from whole blood through centrifugation at 2,000g for 15 min and then stored at −80°C until use. The autoantibodies were studied using a LKS Composite Block Autoantibodies method of indirect immunofluorescence technique, where diluted patient samples and controls are incubated with the Rat liver/kidney/stomach composite blocks. Unbound antibodies are washed off and an appropriate Fluorescein labelled anti human IgG conjugate is applied. Unbound conjugate was washed, and the slides were viewed with a fluorescence/LED microscope. Positive reactions were of bright green colour on relevant tissue. The test was performed on the QuantaLyser SOP BIO 1082.

In addition, antinuclear and cytoplasmic antibodies were tested by indirect immunofluorescence on HEp-2 ANA slides (BioRad) using serum dilution 1:40 and control sera for autofluorescence followed by AlexaFluor488 goat anti-human IgG. Images were acquired using the confocal microscope. ANA screen ELISA (Calbiotech catalogue #AN033G), Doublestranded DNA IgG ELISA (Calbiotech, Catalogue #DD037G) and Mitochondrial Autoantibodies ELISA kits (Calbiotech) were used to determine autoantibodies in patients’ serum samples following the manufacturer’s manual.

### Statistical analysis

The animal experiments were appropriately randomised and blindly allocated for avoidance of bias. Mice were genotyped for the mutations, but the outcome of the phenotype was analysed postmortem, by histopathological examinations and assessments of several markers. Histological scoring assessment was carried out by JA, a certified veterinary pathologist who was blind to genotype information and associated immunological or biochemical markers. The eliglustat treatment was administered with mouse chou, each animal was an experimental unit. Each cage was assigned to treatment groups randomly. Genotype information was not accessible to the technician who was involved in monitoring of mice to allow objective assessment. All offspring received the same amount of inducing agent following the laboratory protocol. Sphingolipid concentrations in tissues were analysed in each experimental animal and the operator was blind to genotypes. Both sexes, males and females were used.

The non-parametric Mann-Whitney test and unpaired two-tailed Student’s tests were used for between groups comparisons. For multiple comparisons one-way ANOVA or Kruskal-Wallis tests were employed. Survival was assessed using Log-Rank test and Kaplan-Meier curve.

## Supplemental material

Figure S1. Glycosylsphingolipids (GlcCer, GlcSph and GlcChol) in tissues of untreated and eliglustat-treated GD mice.

Figure S2. Glycosylsphingolipids (GlcCer, GlcSph and GlcChol) in tissues of untreated and eliglustat-treated Cd1d1-/-d2-/-GCflox/flox Cre+ mice.

Figure S3. Glycosylsphingolipids (GlcCer, GlcSph and GlcChol in serum of untreated and eliglustat-treated Gaucher and Cd1d1-/-d2-/-GCflox/flox Cre+ mice.

Figure S4. Systemic inflammation in a chimeric F1 mixed 129sv/B6 background Gaucher mice. Representative haematoxylin & eosin-stained sections of liver, lung, kidney, spleen and small intestine of Gaucher mice.

Table S1. Mice in the F2 generation.

Table S2 Prevalence of autoantibodies in sera of patients with Gaucher disease (total n=201).

### Human ethics

Anonymised sera from a consortium of UK patients with an enzymatic and genetically confirmed Gaucher disease were collected as part of the Gaucherite study and stored with curation at an approved facility in the Department of Pharmacology University of Oxford, UK according to the principles of the Declaration of Helsinki (2013) (D’Amore et al., 2021). Ethical approval was given by the national ethics committee (14/EE/1168) under the purview of the NHS Research and Development (R&D) in the United Kingdom and from the local R&D boards at each of the eight participating specialist clinical centres. Informed consent was obtained from all subjects at the time of enrolment into GAUCHERITE. The research cohort is registered with ClinicalTrials.gov, NCT code, 03240653.

## Data Availability

All data are presented in the figures and supplementary materials.

## Author Contributions

E.V.P. and T.M.C. conceived of the initial project. E.V.P. designed, supervised and carried out experiments, analysed data and wrote the manuscript. J.A. carried out histology review and scoring tissues. G.D.L.F. carried out lipid analysis, A.J.M.F.G. supervised the mass spectrometry analysis, T.M.C. - project administration, editing of the manuscript, All authors reviewed the manuscript.

## Acknowledgments

We thank Professor Frances M Platt and Dr Kerri Wallom, Department of Pharmacology, University of Oxford, UK for making curated Gaucherite archive sera available for autoantibody studies. We thank Dr Aimee Donald for sharing clinical data on the patients with liver disease as part of the Gaucherite consortium.

We are grateful to Drs Bing Wang, Can Kayatekin and Pablo Sardi of Rare and Neurologic Disease Research, Sanofi R&D, Framingham USA for determining glucosylceramide acyl chain lengths by lc-ms and providing the raw data. Susan Wang for assistance with genotyping, and sample collection.

We thank the Human Research Tissue Bank and Biomedical Research Centre for processing mouse tissues and standard histopathological staining. We thank University Biomedical Services facility at the Anne-McLaren Building for technical support with animal experiments and tissue collections.

## References

Arends, M., L. van Dussen, M. Biegstraaten, and Carla. 2013. Malignancies and monoclonal gammopathy in Gaucher disease; a systematic review of the literature. Br J Haematol. 161:832–842. doi:10.1111/bjh.12335.

Ayto, R., and D.A. Hughes. 2013. Gaucher disease and myeloma. Crit Rev Oncog. 18:247–268. doi:10.1615/critrevoncog.2013006061.

Boer, D.E., M. Mirzaian, M.J. Ferraz, K.C. Zwiers, M. V Baks, M.D. Hazeu, R. Ottenhoff, A.R.A. Marques, R. Meijer, J.C.P. Roos, T.M. Cox, R.G. Boot, N. Pannu, H.S. Overkleeft, M. Artola, and J.M. Aerts. 2021. Human glucocerebrosidase mediates formation of xylosyl-cholesterol by β-xylosidase and transxylosidase reactions. J Lipid Res. 62:100018. doi:10.1194/jlr.ra120001043.

Boven, L.A., M. Van Meurs, R.G. Boot, A. Mehta, L. Boon, J.M. Aerts, and J.D. Laman. 2004. Gaucher cells demonstrate a distinct macrophage phenotype and resemble alternatively activated macrophages. Am J Clin Pathol. 122. doi:10.1309/BG5VA8JRDQH1M7HN.

D’Amore, S., K.R. Page, A. Donald, K. Taiyari, Brian, P. Deegan, C.T. Tan, Kenneth, S. Jones, A.C. Mehta, D. Hughes, R. Sharma, R.H. Lachmann, A. Chakrapani, T. Geberhiwot, S. Santra, S. Banka, and T.M. Cox. 2021. In-depth phenotyping for clinical stratification of Gaucher disease. Orphanet J Rare Dis. 16. doi:10.1186/s13023-021-02034-6

Darmoise, A., P. Maschmeyer, and F. Winau. (2010) The Immunological Functions of Saposins. Adv. Immunol. 105: 25–62. doi10.1016/S0065-2776(10)05002-9

Enquist, I.B., E. Nilsson, A. Ooka, J.-E. Månsson, K. Olsson, M. Ehinger, R.O. Brady, J. Richter, and S. Karlsson. 2006. Effective cell and gene therapy in a murine model of Gaucher disease. Proc Natl Acad Sci U S A. 103:13819–24. doi:10.1073/pnas.0606016103.

Ersek, A., K. Xu, A. Antonopoulos, T.D. Butters, A.E. Santo, Y. Vattakuzhi, L.M. Williams, K. Goudevenou, L. Danks, A. Freidin, E. Spanoudakis, S. Parry, M. Papaioannou, E. Hatjiharissi, A. Chaidos, D.S. Alonzi, G. Twigg, M. Hu, R.A. Dwek, S.M. Haslam, I. Roberts, A. Dell, A. Rahemtulla, N.J. Horwood, and A. Karadimitris. 2015. Glycosphingolipid synthesis inhibition limits osteoclast activation and myeloma bone disease. J Clin Invest. 125:2279–92. doi:10.1172/JCI59987.

de Fost, M., T.A. Out, F.A. de Wilde, E.P.M. Tjin, S.T. Pals, M.H.J. van Oers, R.G. Boot, J.F.M.G. Aerts, M. Maas, S. vom Dahl, and C.E.M. Hollak. 2008. Immunoglobulin and free light chain abnormalities in Gaucher disease type I: data from an adult cohort of 63 patients and review of the literature. Ann Hematol. 87:439–449. doi:10.1007/s00277-008-0441-8.

de Fost, M., S. vom Dahl, G.J. Weverling, N. Brill, S. Brett, D. Häussinger, and C.E.M. Hollak. 2006. Increased incidence of cancer in adult Gaucher disease in Western Europe. Blood Cells Mol Dis. 36:53–58. doi:10.1016/j.bcmd.2005.08.004.

Georgiev, H., I. Ravens, C. Benarafa, R. Förster, and G. Bernhardt. 2016. Distinct gene expression patterns correlate with developmental and functional traits of iNKT subsets. Nat Commun. 7:13116. doi:10.1038/ncomms13116.

Ida, H., O.M. Rennert., T. Ito., K. Maekawa,., and Y. Eto. 1996. Clinical and genetic studies of five fatal cases of Japanese Gaucher disease type 1. Pediatrics International. 38:233–236. doi:10.1111/j.1442-200x.1996.tb03476.x.

James, S.P., F.W. Stromeyer, C. Chang, and J.A. Barranger. 1981. LIver abnormalities in patients with Gaucher’s disease. Gastroenterology. 80:126–133.

Kaplan, M.M. 2004. Novosphingobium aromaticivorans: a potential initiator of primary biliary cirrhosis. Am J Gastroenterol. 99:2147–9. doi:10.1111/j.1572-0241.2004.41121.x.

Kyle, R.A., D.R. Larson, T.M. Therneau, A. Dispenzieri, S. Kumar, J.R. Cerhan, and S.V. Rajkumar. 2018. Long-Term Follow-up of Monoclonal Gammopathy of Undetermined Significance. New England Journal of Medicine. 378:241–249. doi:10.1056/nejmoa1709974.

Lachmann, R.H. 2000. Massive hepatic fibrosis in Gaucher’s disease: clinico-pathological and radiological features. QJM. 93:237–244. doi:10.1093/qjmed/93.4.237.

Lelieveld, L.T., M. Mirzaian, C.J. Kuo, M. Artola, M.J. Ferraz, R.E.A. Peter, H. Akiyama, P. Greimel, H.S. Overkleeft, R.G. Boot, A.H. Meijer, and Johannes. 2019. Role of μ-glucosidase 2 in aberrant glycosphingolipid metabolism: model of glucocerebrosidase deficiency in zebrafish. J Lipid Res. 60:1851–1867. doi:10.1194/jlr.ra119000154.

Leng, H., A.K. Simon, and N.J. Horwood. 2024. Blocking glycosphingolipid production alters autophagy in osteoclasts and improves myeloma bone disease. Autophagy. 20:930–932. doi:10.1080/15548627.2023.2208931.

Leng, H., H. Zhang, L. Li, S. Zhang, Y. Wang, S.J. Chavda, D. Galas-Filipowicz, H. Lou, A. Ersek, E. V Morris, E. Sezgin, Y.-H. Lee, Y. Li, A.V. Lechuga-Vieco, M. Tian, J.-Q. Mi, K. Yong, Q. Zhong, C.M. Edwards, A.K. Simon, and N.J. Horwood. 2022. Modulating glycosphingolipid metabolism and autophagy improves outcomes in pre-clinical models of myeloma bone disease. Nat Commun. 13:7868. doi:10.1038/s41467-022-35358-3.

Manning, L.S., and N.S. Radin. 1999. Effects of the glucolipid synthase inhibitor, P4, on functional and phenotypic parameters of murine myeloma cells. Br J Cancer. 81:952–8. doi:10.1038/sj.bjc.6690792.

Marques, A.R.A., M. Mirzaian, H. Akiyama, P. Wisse, M.J. Ferraz, P. Gaspar, K. Ghauharali-van der Vlugt, R. Meijer, P. Giraldo, P. Alfonso, P. Irún, M. Dahl, S. Karlsson, E. V Pavlova, T.M. Cox, S. Scheij, M. Verhoek, R. Ottenhoff, C.P.A.A. van Roomen, N.S. Pannu, M. van Eijk, N. Dekker, R.G. Boot, H.S. Overkleeft, E. Blommaart, Y. Hirabayashi, and J.M. Aerts. 2016. Glucosylated cholesterol in mammalian cells and tissues: formation and degradation by multiple cellular β-glucosidases. J Lipid Res. 57:451–63. doi:10.1194/jlr.M064923.

Mattner, J., P.B. Savage, P. Leung, S.S. Oertelt, V. Wang, O. Trivedi, S.T. Scanlon, K. Pendem, L. Teyton, J. Hart, W.M. Ridgway, L.S. Wicker, M.E. Gershwin, and A. Bendelac. 2008. Liver autoimmunity triggered by microbial activation of natural killer T cells. Cell Host Microbe. 3:304–15. doi:10.1016/j.chom.2008.03.009.

McEachern, K. A., J. Fung., S. Komarnitsky., C.S. Siegel., W.L. Chuang., E. Hutto., J.A. Shayman., G.A. Grabowski., J.M. Aerts., S.H. Cheng., D.P. Copeland, and J Marshall. 2007. A specific and potent inhibitor of glucosylceramide synthase for substrate inhibition therapy of Gaucher disease. Mol Genet Metab. 91:259–267. doi10.1016/j.ymgme.2007.04.001

Mistry, P.K., T.H. Taddei, S. Vom Dahl, and B.E. Rosenbloom. 2013. Gaucher Disease and Malignancy: A Model for Cancer Pathogenesis in an Inborn Error of Metabolism. Crit Rev Oncog. 18:235–246. doi:10.1615/critrevoncog.2013006145.

Mistry, P. K., M. Balwani., J. Charrow., J. Lorber., C. Niederau., J.L. Carwile., A. Oliveira-Dos-Santos., M.G. Perichon., S. Uslu Cil, and P.S. Kishnani.2024. Long-term effectiveness of eliglustat treatment: A real-world analysis from the International Collaborative Gaucher Group Gaucher Registry. Am J Hematol. 99: 1500–1510. doi10.1002/ajh.27347

Nair, S., C.S. Boddupalli, R. Verma, J. Liu, R. Yang, G.M. Pastores, P.K. Mistry, and M. V Dhodapkar. 2015. Type II NKT-TFH cells against Gaucher lipids regulate B-cell immunity and inflammation. Blood. 125:1256–1271. doi:10.1182/blood-2014-09-600270.

Nair, S., A.R. Branagan, J. Liu, C.S. Boddupalli, P.K. Mistry, and M. V Dhodapkar. 2016. Clonal Immunoglobulin against Lysolipids in the Origin of Myeloma. New England Journal of Medicine. 374:555–561. doi:10.1056/nejmoa1508808

Nair, S., J. Sng, C.S. Boddupalli, A. Seckinger, M. Chesi, M. Fulciniti, L. Zhang, N. Rauniyar, M. Lopez, N. Neparidze, T. Parker, N.C. Munshi, R. Sexton, B. Barlogie, R. Orlowski, L. Bergsagel, D. Hose, R.A. Flavell, P.K. Mistry, E. Meffre, and M. V Dhodapkar. 2018. Antigen-mediated regulation in monoclonal gammopathies and myeloma. JCI Insight. 3. doi:10.1172/jci.insight.98259.

Nascimbeni, F., E. Cassinerio, A.D. Salda, I. Motta, S. Bursi, S. Donatiello, V. Spina, M.D. Cappellini, and F. Carubbi. 2018. Prevalence and predictors of liver fibrosis evaluated by vibration controlled transient elastography in type 1 Gaucher disease. Mol Genet Metab. 125:64–72. doi:10.1016/j.ymgme.2018.08.004.

Nguyen, Y., J. Stirnemann., F. Lautredoux., B. Cador., M. Bengherbia., K. Yousfi., D. Hamroun., L Astudillo., T. Billette de Villemeur., A. Brassier., F. Camou., F. Dalbies., D. Dobbelaere., F. Gaches., V. Leguy-Seguin., A. Masseau., Y.M. Pers., S. Pichard., C. Serratrice., M.G. Berger., B. Fantin and N.Belmatoug. 2020. On Behalf Of The French Evaluation Of Gaucher Disease Treatment Committee (2020). Immunoglobulin Abnormalities in Gaucher Disease: an Analysis of 278 Patients Included in the French Gaucher Disease Registry. Internat J Mol Sci, 21(4), 1247. doi.org/10.3390/ijms21041247

Park, J.-W., W.-J. Park, and A.H. Futerman. 2014. Ceramide synthases as potential targets for therapeutic intervention in human diseases. Biochim Biophys Acta. 1841:671–81. doi:10.1016/j.bbalip.2013.08.019.

Patel, S.C., G.L. Davis, and J.A. Barranger. 1986. Gaucher’s disease in a patient with chronic active hepatitis. Am J Med. 80:523–525. doi:10.1016/0002-9343(86)90734-5.

Pavlova, E. V, S.Z. Wang, J. Archer, N. Dekker, J. Aerts, S. Karlsson, and T.M. Cox. 2013a. B cell lymphoma and myeloma in murine Gaucher’s disease. J Pathol. 231:88–97. doi:10.1002/path.4227.

Pavlova, E. V., J. Archer., S. Wang., N. Dekker., J.M. Aerts., S. Karlsson., and T.M. Cox. 2015. Inhibition of UDP-glucosylceramide synthase in mice prevents Gaucher disease-associated B-cell malignancy. Journal Pathol. 235:113–124. doi10.1002/path.4452

Regenboog, M., L. van Dussen, J. Verheij, N.J. Weinreb, D. Santosa, S. vom Dahl, D. Häussinger, M.N. Müller, A. Canbay, M. Rigoldi, A. Piperno, T. Dinur, A. Zimran, P.K. Mistry, K.Y. Salah, N. Belmatoug, D.J. Kuter, and Carla. 2018. Hepatocellular carcinoma in Gaucher disease: an international case series. J Inherit Metab Dis. 41:819–827. doi:10.1007/s10545-018-0142-y.

Rosenbloom, B. E., and N. J. Weinreb. 2013. Gaucher disease: a comprehensive review. Critical reviews in oncogenesis, 18: 163–175. doi10.1615/critrevoncog.2013006060

Rosenbloom, B.E., M.D. Cappellini, N.J. Weinreb, M. Dragosky, S. Revel-Vilk, J.L. Batista, D. Sekulic, and P.K. Mistry. 2022a. Cancer risk and gammopathies in 2123 adults with Gaucher disease type 1 in the International Gaucher Group Gaucher Registry. Am J Hematol. 97:1337–1347. doi:10.1002/ajh.26675.

Serratrice, C., N. Bensalah., G. Penaranda., N. Bardin., N. Belmatoug., A. Masseau., C. Rose., O. Lidove., F. Camou., F. Maillot., V. Leguy., N. Magy-Bertrand., I. Marie., P. Cherin., M. Bengherbia., S. Carballo., J. Boucraut., J. Serratrice., M. Berger., and D Verrot. 2018. Prevalence of autoantibodies in the course of Gaucher disease type 1: A multicenter study comparing Gaucher disease patients to healthy subjects. Joint bone spine, 85(1), 71–77. doi10.1016/j.jbspin.2016.12.002

Shoenfeld, Y., and E. Mozes. 1990. Pathogenic idiotypes of autoantibodies in autoimmunity: lessons from new experimental models of SLE. The FASEB Journal. 4:2646–2651. doi:10.1096/fasebj.4.9.2140806.

Shoenfeld, Y., A. Beresovski, D. Zharhary, Y. Tomer, M. Swissa, E. Sela, A. Zimran, S. Zevin, B. Gilburd, and M. Blank. 1995. Natural autoantibodies in sera of patients with Gaucher’s disease. J Clin Immunol. 15:363–372. doi:10.1007/bf01541326.

Sigmon, J.S., M.W. Blanchard, R.S. Baric, T.A. Bell, J. Brennan, G.A. Brockmann, A.W. Burks, J.M. Calabrese, K.M. Caron, R.E. Cheney, D. Ciavatta, F. Conlon, D.B. Darr, J. Faber, C. Franklin, T.R. Gershon, L. Gralinski, B. Gu, C.H. Gaines, R.S. Hagan, E.G. Heimsath, M.T. Heise, P. Hock, F. Ideraabdullah, J.C. Jennette, T. Kafri, A. Kashfeen, M. Kulis, V. Kumar, C. Linnertz, A. Livraghi-Butrico, K.C.K. Lloyd, C. Lutz, R.M. Lynch, T. Magnuson, G.K. Matsushima, R. McMullan, D.R. Miller, K.L. Mohlke, S.S. Moy, C.E.Y. Murphy, M. Najarian, L. O’Brien, A.A. Palmer, B.D. Philpot, S.H. Randell, L. Reinholdt, Y. Ren, S. Rockwood, A.R. Rogala, A. Saraswatula, C.M. Sassetti, J.C. Schisler, S.A. Schoenrock, G.D. Shaw, J.R. Shorter, C.M. Smith, C.L. St Pierre, L.M. Tarantino, D.W. Threadgill, W. Valdar, B.J. Vilen, K. Wardwell, J.K. Whitmire, L. Williams, M.J. Zylka, M.T. Ferris, L. McMillan, and F.P. Manuel de Villena. 2020. Content and Performance of the MiniMUGA Genotyping Array: A New Tool To Improve Rigor and Reproducibility in Mouse Research. Genetics. 216:905–930. doi:10.1534/genetics.120.303596.

Smiley, S.T., M.H. Kaplan, and M.J. Grusby. 1997. Immunoglobulin E production in the absence of interleukin-4-secreting CD1-dependent cells. Science. 275:977–9. doi:10.1126/science.275.5302.977.

Stirnemann, J., N. Belmatoug., F. Camou., C. Serratrice., R. Froissart., C. Caillaud., T. Levade., L. Astudillo., J. Serratrice., A. Brassier., C. Rose., T. Billette de Villemeur, and M. G. Berger. 2017. A Review of Gaucher Disease Pathophysiology, Clinical Presentation and Treatments. Internat. J Mol Sci. 18(2), 441. doi.org/10.3390/ijms18020441

Ullman-Culleré, M.H., and C.J. Foltz. 1999. Body condition scoring: a rapid and accurate method for assessing health status in mice. Lab Anim Sci. 49:319–23.

Weinreb, N.J., D.S. Barbouth, and R.E. Lee. 2018. Causes of death in 184 patients with type 1 Gaucher disease from the United States who were never treated with enzyme replacement therapy. Blood Cells Mol Dis. 68:211–217. doi:10.1016/j.bcmd.2016.10.002.

Weinreb, N.J., and R.E. Lee. 2013. Causes of death due to hematological and non-hematological cancers in 57 US patients with type 1 Gaucher Disease who were never treated with enzyme replacement therapy. Crit Rev Oncog. 18:177–195. doi:10.1615/critrevoncog.2013005921.

Zaccariotto, E., M.B. Cachón-González, B. Wang, S. Lim, B. Hirth, H. Park, M. Fezoui, S.P. Sardi, P. Mason, R.H. Barker, and T.M. Cox. 2022. A novel brain-penetrant oral UGT8 inhibitor decreases in vivo galactosphingolipid biosynthesis in murine Krabbe disease. Biomed Pharmacother. 149:112808. doi:10.1016/j.biopha.2022.112808.

