## Supplementary material for "The Role of Glycosphingolipids in Autoimmune Manifestations and Myeloma in Gaucher Disease"

**Figure S1.** Glycosylsphingolipids (GlcCer, GlcSph and GlcChol) in tissues of untreated and eliglustat-treated GD mice.

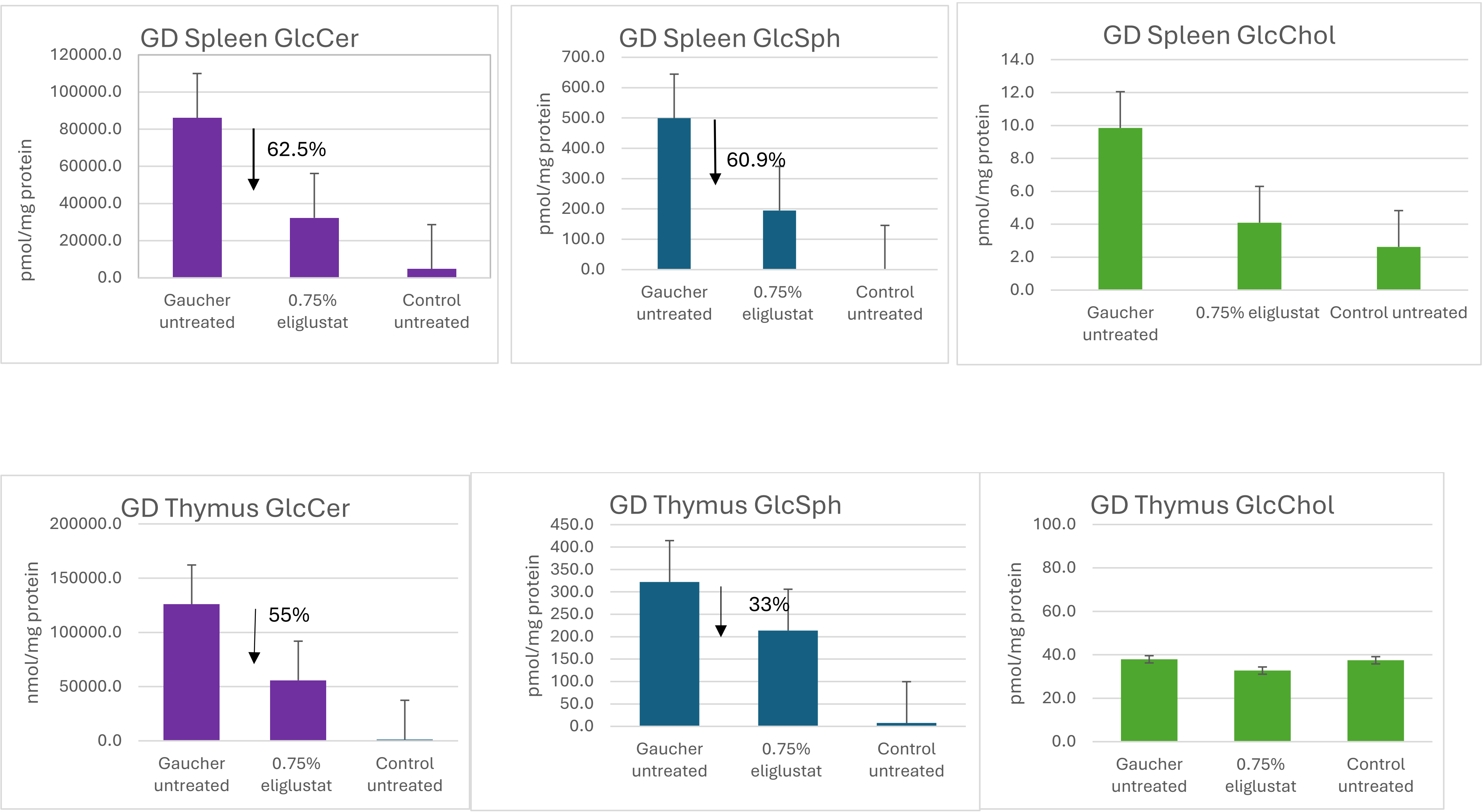

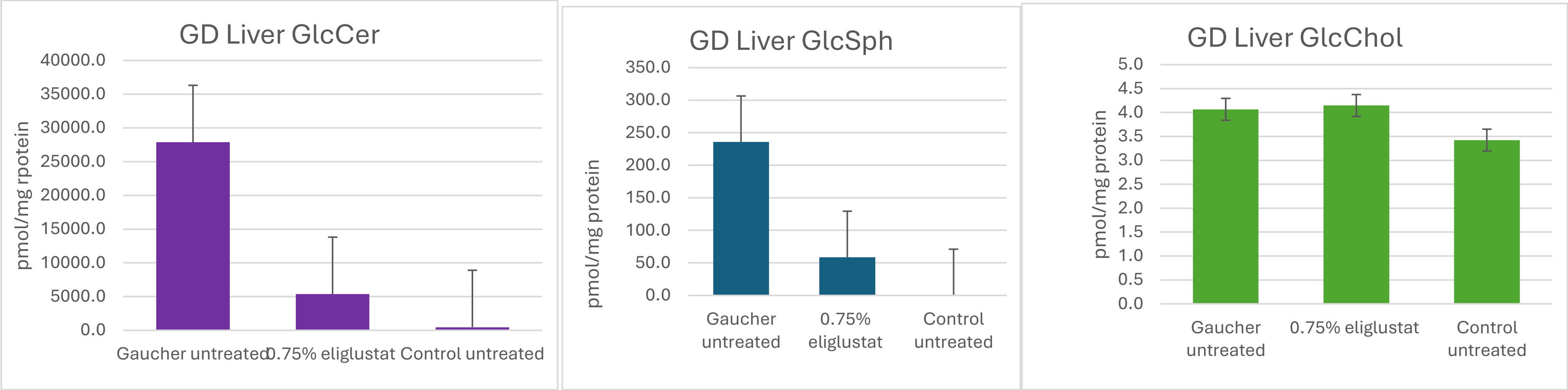


**Figure S2.** Glycosylsphingolipids (GlcCer, GlcSph and GlcChol in tissues of untreated and eliglustat-treated Cd1d1$^{-/-}$d2$^{-/-}$ GC$^{flox/flox}$ Cre+ mice.

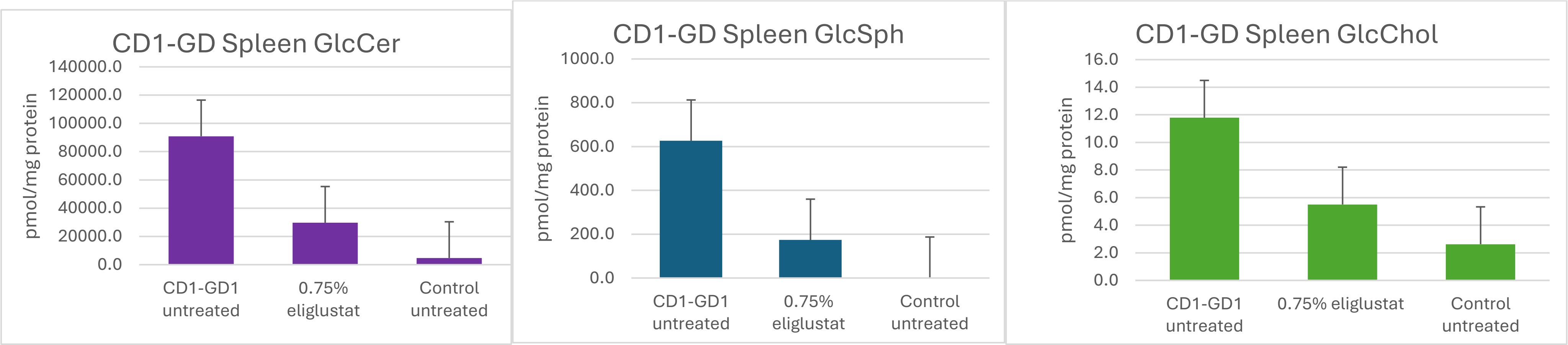

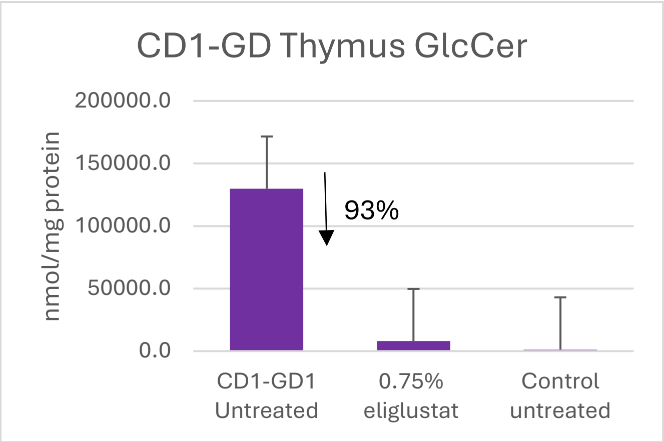
CD1-GD Thymus GlcCer
nmol/mg protein
200000.0
150000.0
100000.0
50000.0
0.0
93%
CD1-GD1 Untreated
0.75% eliglustat
Control untreated

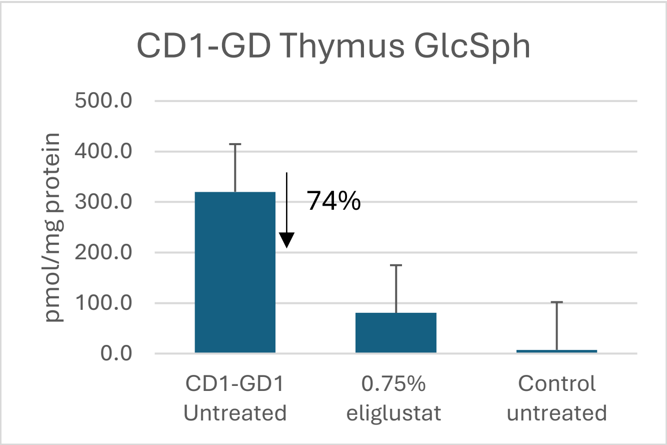
CD1-GD Thymus GlcSph
pmol/mg protein
500.0
400.0
300.0
200.0
100.0
0.0
74%
CD1-GD1 Untreated
0.75% eliglustat
Control untreated

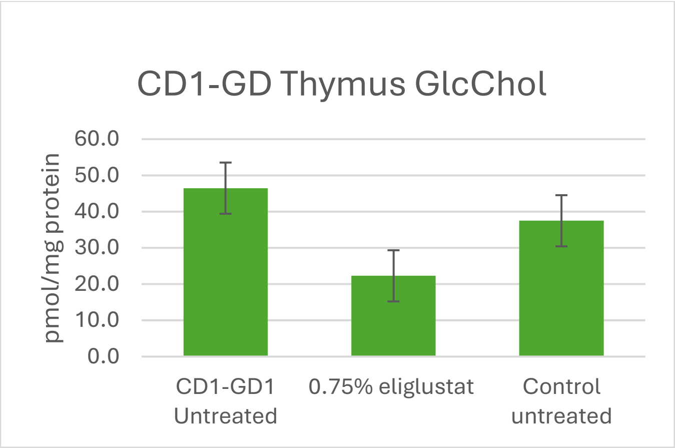
CD1-GD Thymus GlcChol
pmol/mg protein
60.0
50.0
40.0
30.0
20.0
10.0
0.0
CD1-GD1 Untreated
0.75% eliglustat
Control untreated

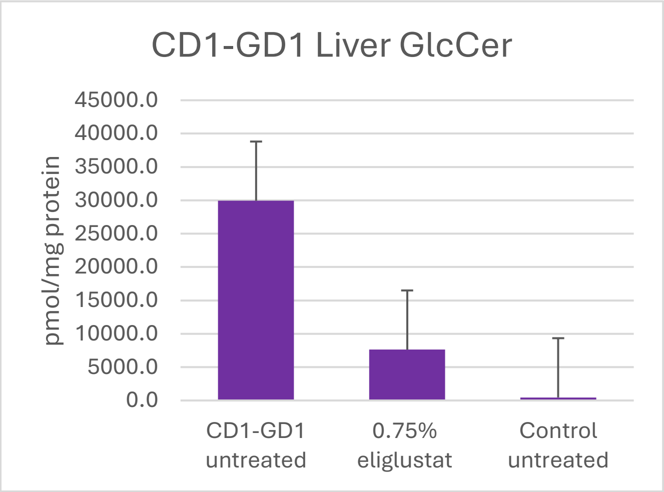
CD1-GD1 Liver GlcCer
pmol/mg protein
45000.0
40000.0
35000.0
30000.0
25000.0
20000.0
15000.0
10000.0
5000.0
0.0
CD1-GD1 untreated
0.75% eliglustat
Control untreated

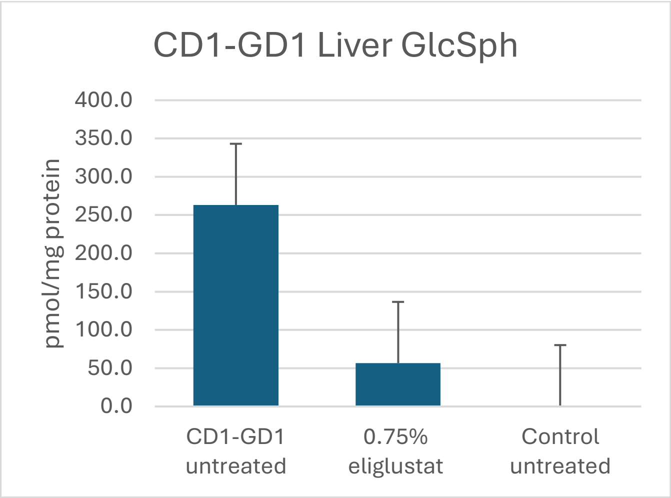
CD1-GD1 Liver GlcSph
pmol/mg protein
400.0
350.0
300.0
250.0
200.0
150.0
100.0
50.0
0.0
CD1-GD1 untreated
0.75% eliglustat
Control untreated

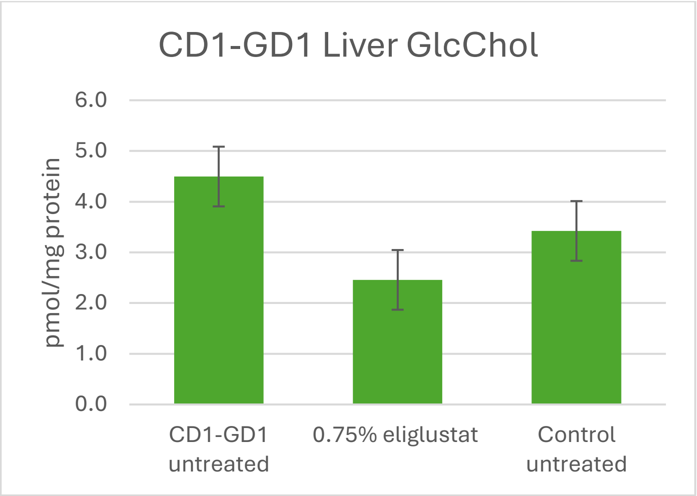
CD1-GD1 Liver GlcChol
pmol/mg protein
6.0
5.0
4.0
3.0
2.0
1.0
0.0
CD1-GD1 untreated
0.75% eliglustat
Control untreated

**Figure S3.** Glycosylsphingolipids (GlcCer, GlcSph and GlcChol in serum of untreated and eliglustat-treated Gaucher and Cd1d1$^{-/-}$d2$^{-/-}$ GC$^{flox/flox}$ Cre+ mice.

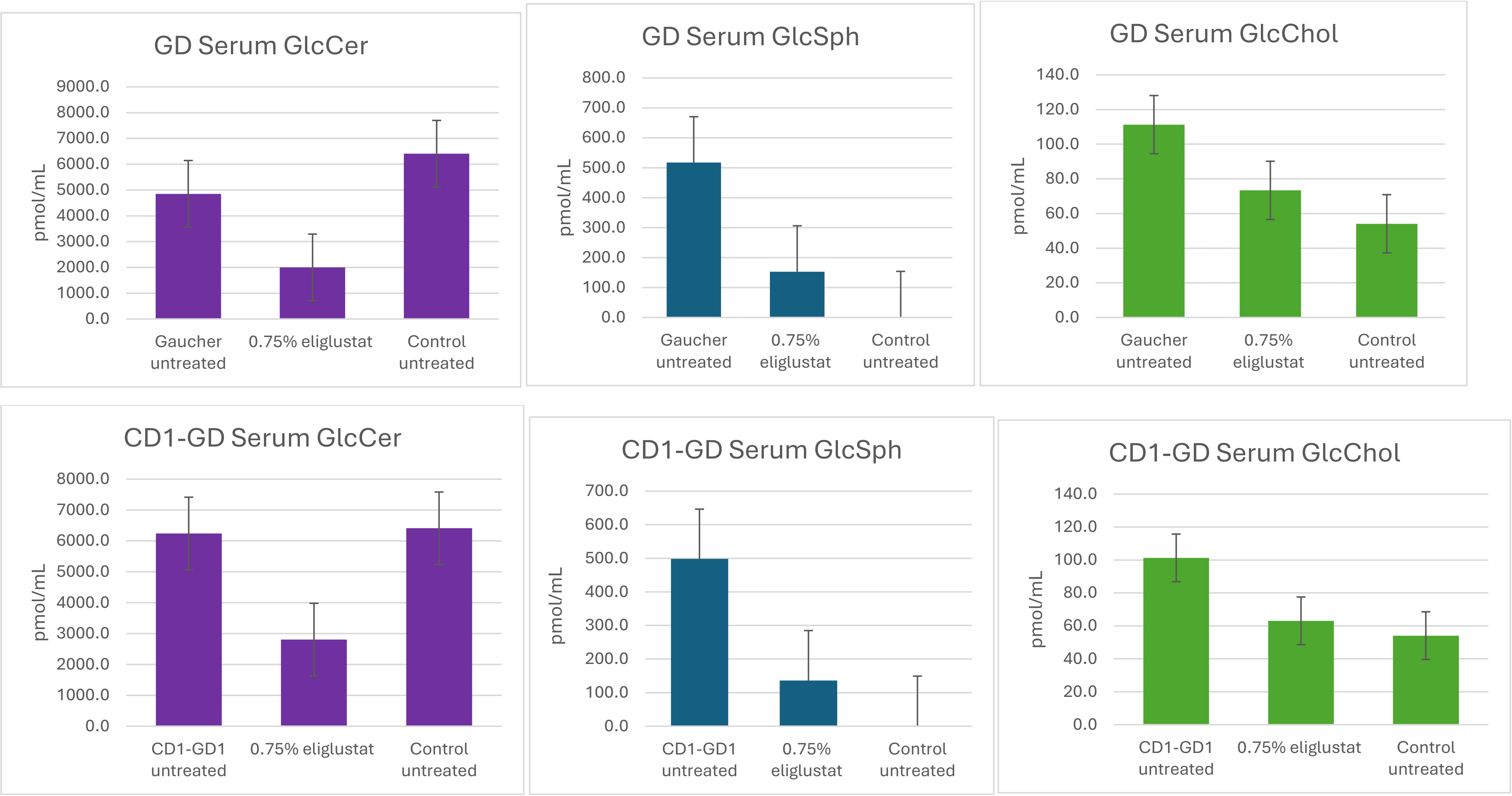

Table S1. Mice in the F2 generation

| Genotype | Number of mice | Nature of related gametes |
|---|---|---|
| $CD1^{+/-}$ $GC^{flox/wt}$ | 29 | Nonrecombinant |
| $CD1^{+/+}$ $GC^{flox/flox}$ | 12 | Nonrecombinant |
| $CD1^{-/-}$ $GC^{wt/wt}$ | 18 | Nonrecombinant |
| $CD1^{+/-}$ $GC^{flox/flox}$ | 1 | Recombinant |
| $CD1^{+/-}$ $GC^{wt/wt}$ | 4 | Recombinant |
| $CD1^{+/+}$ $GC^{flox/wt}$ | 2 | Recombinant |

Table S2. Prevalence of autoantibodies in sera of patients with Gaucher disease (total n=201).

| | N autoantibodies positive | N autoantibodies weakly positive | Total |
|---|---|---|---|
| Females | 7 | 12 | 19 |
| Males | 7 (+1 unknown) | 7 | 14 |
| Total | 15 | 19 | 34 |

Figure S4. Systemic inflammation in a chimeric F1 mixed 129sv/B6 background Gaucher mice. Representative haematoxylin & eosin stained sections of liver, lung, kidney, spleen and small intestine of Gaucher mice.

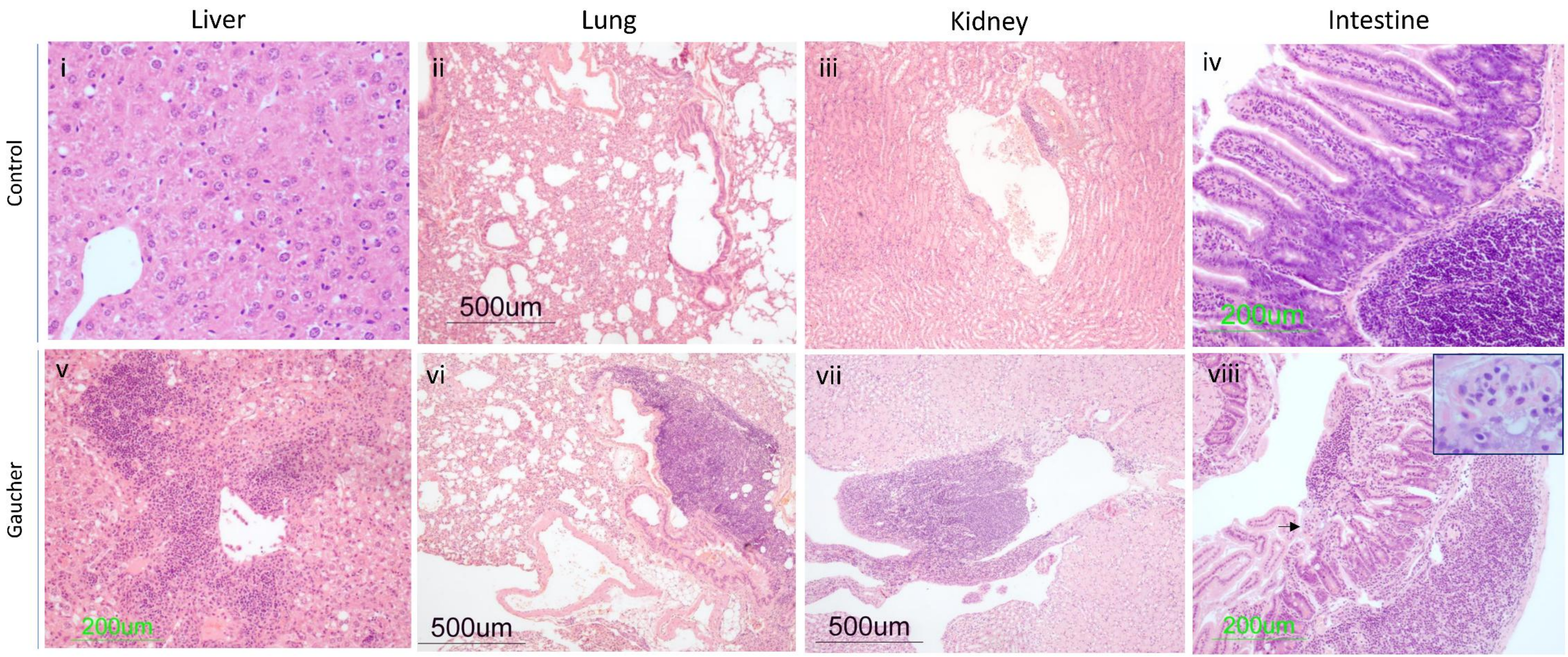
